# The ERK5/NF-κB signaling pathway targets endometrial cancer proliferation and survival

**DOI:** 10.1101/2022.05.31.494135

**Authors:** Nora Diéguez-Martínez, Sergio Espinosa-Gil, Guillermo Yoldi, Elisabet Megías-Roda, Idoia Bolinaga-Ayala, Maria Viñas-Casas, Inés Domingo-Ortí, Héctor Pérez-Montoyo, Jose R Bayascas, Eva Colas, Xavier Dolcet, Jose M Lizcano

## Abstract

Endometrial cancer (EC) is the most common type of gynaecologic cancer in women of developed countries. Despite surgery combined with chemo-/radiotherapy regimens, overall survival of patients with high-risk EC tumors is poor, indicating a need for novel molecular therapies. The MEK5-ERK5 pathway is activated in response to growth factors and to different forms of stress, including oxidative stress and cytokines. Previous evidence support a role for the MEK5-ERK5 pathway in the pathology of several cancers. We have investigated the role of ERK5 in EC. In silico analysis of the PanCancer Atlas dataset showed alterations in components of the MEK5-ERK5 pathway in 48% of EC patients. Here, we show that ERK5 inhibition decreased EGF-induced EC cell proliferation, and that depletion of MEK5 resulted in EC impaired proliferation and reduced tumor growth capacity in nude mice. Pharmacologic or genetic silencing of ERK5 impaired NF-kB pathway in EC cells and xenografts. Furthermore, we found a positive correlation between ERK5 and p65/RELA protein levels in human EC tumor samples. Mechanistically, impairment of ERK5 resulted in downregulation of NEMO/IKKγ expression, leading to impaired p65/RELA activity and to apoptosis in EC cells and xenografts, which was rescued by NEMO/IKKγ overexpression. Notably, ERK5 inhibition, MEK5 depletion or NF-kB inhibition sensitized EC cells to standard EC chemotherapy (paclitaxel/carboplatin) toxicity, whereas ERK5 inhibition synergized with paclitaxel to reduce tumor xenograft growth in mice. Together, our results suggest that the ERK5-NEMO-NF-κB pathway mediates EC cell proliferation and survival. We propose the ERK5/NF-κB axis as new target for EC treatment.

## Introduction

Endometrial cancer (EC) EC is the most common type of gynaecological cancer and the fourth most frequent neoplasia in women of developed countries [1]. EC originates from the uterine endometrial epithelium and is histological classified into two different histological subtypes [2]. Endometrioid EC accounts for the majority of ECs. These tumors express estrogen and/or progesterone receptors, which are used as targets for hormonal therapy. Non-endometrioid EC (NEEC) represents ∼20% of cases and comprises serous and clear cell carcinomas, among others. Non-endometrioid EC tumors tend to be estrogen-independent and show enhanced aggressiveness and worse prognosis than endometrioid ECs [2].

Advanced and non-hormone-dependent ECs are currently treated with systemic cytotoxic therapy with carboplatin/paclitaxel [3]. However, its clinical effectiveness is poor in advanced, recurrent or metastasic EC tumors, indicating a need for novel molecular therapies. Since EC tumors frequently concur with activating mutations of *PI3KCA* or *AKT*, the PI3K-AKT-mTOR pathway has been proposed for EC targeted therapies. Unfortunately, EC tumors frequently develop resistance to PI3K, Akt, or mTOR inhibitors, mainly due to hyperactivation of the Ras-MEK1 pathway [4]. Hence, there is an urgent need for new targeted therapies to improve treatment of high-risk EC.

The extracellular regulated kinase 5 (ERK5) is the most recent discovered member of the conventional MAP kinase family. ERK5 is activated by its unique MAP2K MEK5, in response to both mitogens and different forms of stress, including oxidative stress and cytokines. These effectors, which lead to the activation of the MEK5 upstream activators MAP3K2 and MAP3K3 [5]. ERK5 is involved in the proliferation, survival, apoptosis, and differentiation of different cell types [6]. Recently, ERK5 was shown to play an important role in the tumorigenesis, proliferation and survival of several cancer types (7-8). Because ERK5 inhibition or silencing compromises the viability of numerous cancer cell lines and tumor xenografts, ERK5 is emerging as a new target for molecular anticancer therapies [9].

In this work, we have investigated for the first time the role of the MEK5-ERK5 pathway in EC. We also report the preclinical development of the small-molecule JWG-071, a selective ERK5 inhibitor with a good pharmacokinetic profile. We present evidence supporting ERK5 inhibition as a targeted therapy with great potential to treat EC.

## Materials and methods

### Reagents

JWG-071 was in-house synthesized as previously described [10]. AX-15836, BAY-117082, paclitaxel and carboplatin were obtained from MedChem Express (Monmouth Junction, NJ, USA). Cisplatin, staurosporine and EGF were from Sigma-Aldrich (Germany). Q-VD(OMe)-OPh was from APExBio (Gentaur, Spain).

### Cells and cell culture

AN3CA and ARK1 EC cells and Hela cervical cancer cells were purchased from ATCC (Manassas, VA, USA). Ishikawa cells were from ECACC and were purchase from Sigma. T-large antigen-transformed BAX^-/-^/BAK^-/-^ MEFs were generously provided by Dr Victor Yuste (UAB, Spain). HeLa, ARK1 and MEF cells were maintained in Dulbecco’s modified Eagle’s medium (DMEM ThermoFisher). AN3CA were maintained in DMEM-F12 medium (ThermoFisher). Medium was supplemented with 5% (Ishikawa cells) or 10% (rest of cells) foetal bovine serum (FBS, Gibco) and 1% Penicillin/Streptomycin (Gibco). Cells were maintained at 37°C in a humidified atmosphere containing 5% CO_2_. Cell lines were transfected using Lipofectamine 2000™ (ThermoFisher Scientific, Waltham, MA, USA) as described before [11].

### Cell lysis and immunoblotting

Cells were lysed in ice-cold RIPA buffer supplemented with 1 mM sodium-orthovanadate, 50 mM NaF and 5 mM sodium-pyrophosphate, sonicated and stored at −20°C. Subcellular fractionation was performed following standard procedures, as described before [12]. Proteins were resolved in SDS-PAGE gels and electrotransferred onto a nitrocellulose membrane (Sigma-Aldrich). After incubation with the appropriated primary, detection was performed using horseradish peroxidase-conjugated secondary antibodies and enhanced chemiluminescence reagent (Bio-Rad). Primary antibodies used are given in **Supplementary Table S1**.

### DNA constructs

The pEBG2T vector encoding GST-tagged human ERK5 was previously described [12]. The pCDNA3 vector encoding for SR-IkBα was from J.X. Comella (VHIR Barcelona, Spain)[13]. pGL4-AP1 (containing six copies of an AP1-response element fussed to luciferase gene) and PGLA4.32-NF-κB (five copies of a NF-κB response element) vectors were from Promega (Madison, WI, USA).

### shERK5 lentiviral production

Stable and efficient silencing of endogenous ERK5 protein was achieved using lentivirus. Two pLKO.1 lentiviral vectors encoding for specific shRNA for two human MAPK7 mRNA sequences were used (TRCN0000010262/pLKO.1 and TRCN0000010275/pLKO.1, Sigma). Lentiviral particles were generated in HEK-293 cells by co-transfecting the virion vectors (psPAX2 and pMD2G) and the pLKO.1 ERK5-shRNA vectors. After 4h, the medium was replaced with fresh medium. Forty-eight h post-transfection, the medium containing the viral particles was collected, centrifuged, filtered to remove cell debris, and stored at -80°C until use.

### Gene silencing

Validated siRNAs targeting NEMO were from Sigma (ref. EHU032271) and ThermoFisher (ref. AM51331). Scrambled (control) siRNA was from ThermoFisher. siRNAs were transfected into cells using Lipofectamine-2000, following the manufacturer’s recommended protocol. The vectors encoding two different shRNAs targeting ERK5 were from Sigma (TRCN0000010262/pLKO.1; TRCN0000010275/pLKO.1). Lentiviral production is detailed at Supplementary Materials and Methods.

### Quantitative real-time PCR

Total RNA was isolated from cells using RNeasy kit (Qiagen). cDNA was obtained using iScript™ cDNA Synthesis Kit (Bio-Rad). Real-time quantitative PCR assays were performed using TaqMan Gene Expression Master Mix and the probes human NEMO/IKBKG (Hs00415849_m1) and human GAPDH (Hs03929097_g1) (ThermoFisher Scientific). Amplifications were run in a Bio-Rad CFX96 real-time PCR system, using the following protocol: 50°C for 2 min, 95°C for 10 min, 39 cycles of 95°C for 15 s, 55°C for 1 min. Each value was normalized to GAPDH levels. Relative expression levels were determined using the 2^−ΔΔCt^ method. Real-time PCR was controlled by the Bio-Rad CFX Manager v 3.1 software.

### Generation of MEK5 knockout cells by CRISPR/Cas9

To generate CRISPR/Cas9 MEK5^-/-^ Hela or Ishikawa cells, we used a kit from Santa Cruz technology, using vectors containing sgRNAs (TACTTGCTGTTCCATTACTG and CAGATTCACTTCCAAGCAAT) and the Cas9 endonuclease gene (sc-401688), in combination with a HDR-puromycin resistance vector (sc-401688). Cells were selected with 1 µg/ml puromycin (Sigma). Single colonies were picked up and monitored for MEK5 expression. Clones showing MEK5 null expression were selected.

### Cell viability, clonogenic and proliferation assays

Cell viability was determined by MTT (3-(4,5-dimethyl-2-thiazolyl)-2,5-diphenyl-2H-tetrazolium Blue, Sigma) reduction assay. For clonogenic assays, cells were seeded in a 6 wells-plates, treated for 14 days and stained with 0.5% crystal violet solution. Colonies with >40 cells were counted. Cell proliferation was measured by cell counting assay, using a TC10TM automated cell counter (Bio-Rad, Hercules, CA, USA) using Trypan Blue. Cell viability was also determined using a LIVE/DEAD viability/cytotoxicity kit assay (ThermoFisher), as reported before [11].

### Flow cytometry analysis of apoptosis

Cells were seeded and treated for the indicated times. Cell were then trypsinized, washed sequentially with FBS and binding buffer, and incubated in binding buffer with Annexin V and/or propidium iodide (Invitrogen, Waltham, MA, USA) for 15 min. Samples were analyzed in a Beckman Coulter FC500 flow cytometer. For nuclear staining, cells were fixed with 4% PFA, 0.1% NP40 in PBS, and nuclei were stained with 1 μg/mL Hoechst 33258 (Invitrogen).

### Human EC tumor samples

Patients with no previous therapy (50–80 years) underwent surgery for endometrial carcinoma at Hospital Vall d’Hebron (Barcelona, Spain). The protocol was approved by the Institutional Review Boards, and informed consent was obtained from the patients involved. After surgery, endometrial normal tissue and tumor samples were collected from each patient and stored frozen (−80°C) until analysis. Samples were lysed in RIPA buffer containing protease and phosphatase inhibitor cocktails (Sigma), and stored at -20°C until use.

### Gene reporter Luciferase assay

Cells cultured in 12-well plates were transfected with 450 ng of NF-κB-driven (Stratagene) or AP 1-driven luciferase (Promega) reporter plasmids, and 50 ng Renilla luciferase (Promega). Luciferase activity assay was monitored using the dual luciferase kit (Promega), following the manufacturer’s instructions.

### EC xenografts

Athymic nude female mice (Hsd: Athymic Nude-Foxn1nu, Envigo, Spain) were injected subcutaneously with 4×10^6^ Ishikawa cells into the right flank. When tumors reached 80-100 mm^3^, mice were randomly distributed into groups and administered with the indicated treatments. JWG-071 was administered once a day intraperitoneally at dose of 50 mg/kg (monotherapy) or 30 mg/kg (combo treatment). Paclitaxel was administered twice a week intraperitoneally at dose of 15 mg/Kg. Tumors volumes were measured as (length x width^2^)/2, thrice a week. When tumors reached more than 1000 mm^3^ or body weight decreased 20% of the initial measure, mice were euthanized by cervical dislocation. Tumors were excised from mice, and a fraction of each tumor was homogenized in RIPA buffer containing protease inhibitor cocktail (Sigma), sonicated, centrifuged to remove cell debris, and stored at -20°C until use. All procedures involving animals were performed with approval of the UAB Animal Experimentation Committee, according to Spanish official regulations. Individual tumor growth curves are given in **Supplementary Figure 12**.

### Immunohistochemistry

Samples from tumor xenografts were dissected, formalin-fixed and paraffin-embedded as described previously [14]. Three μm-thickness sections were stained with haematoxylin and analyzed by immunohistochemistry using standard protocols. Briefly, deparaffinized samples were subjected to heat-induced antigen retrieval in citrate buffer. After blocking endogenous peroxidase with 3% H_2_O_2_, and reducing unspecific binding with and 5% goat serum and 0,1% Triton X-100, slices were then incubated with anti-cleaved caspase 3 antibody or anti-Ki67 antibody (Cell Signalling) and further processed with secondary antibody (LSAB2 horseradish peroxidase kit; Dako, Copenhagen, Denmark).

### Statistical analysis

All in vitro data was assessed using one-way ANOVA followed by Bonferroni multiple comparison test or two-tailed Student’s t-test. Tumor volumes of mice were compared using two-way ANOVA followed by Bonferroni, except for combinatorial experiment (two-way ANOVA Tukey’s test). Statistical significance between the groups was assessed with the log-rank test (GraphPad software, San Diego, CA, USA). Data in Figures are presented as mean ± SD, unless otherwise indicated. Statistical significance was set at p < 0.05. Synergism analysis was performed using the Compusyn software [15].

## Results

### ERK5 Signaling pathway is altered in EC

To investigate the impact of the MEK5-ERK5 signaling pathway in EC, we performed an in silico data analysis of the components of this pathway. Gene expression data mining from the cBioPortal for Cancer Genomics (Uterine corpus endometrial carcinoma TCGA, PanCancer Atlas) showed alterations (gene copy number, mutations or mRNA alterations) in all components of ERK5 signaling, including the upstream kinase activators *MAP3K2* and *MAP3K3*, the MEK5 and ERK5 genes (*MAP2K5* and *MAPK7*, respectively), and the ERK5 substrates *MEF2A, MEF2B, MEF2C*, and *MEF2D* transcription factors [16]. Of note, 48% of human EC patients (250 out of 517) showed alterations of components of ERK5 signaling, mostly associated to gene amplification and high mRNA levels (33%, 171 out of 517) (**Fig. 1A**). Interestingly, EC is the human cancer that showed the higher percentage of cases wherein ERK5, MEK5, MAP3K2 and MAP3K3 are mutated (**Supplementary Fig. 1**). Finally, patients with high levels of ERK5 mRNA expression showed lower overall survival (p = 0.000087) and reduced progression-free survival (0.00094) (**Fig. 1B**). Taken together, these data suggest a role of ERK5 signaling in EC.

**Figure 1.**
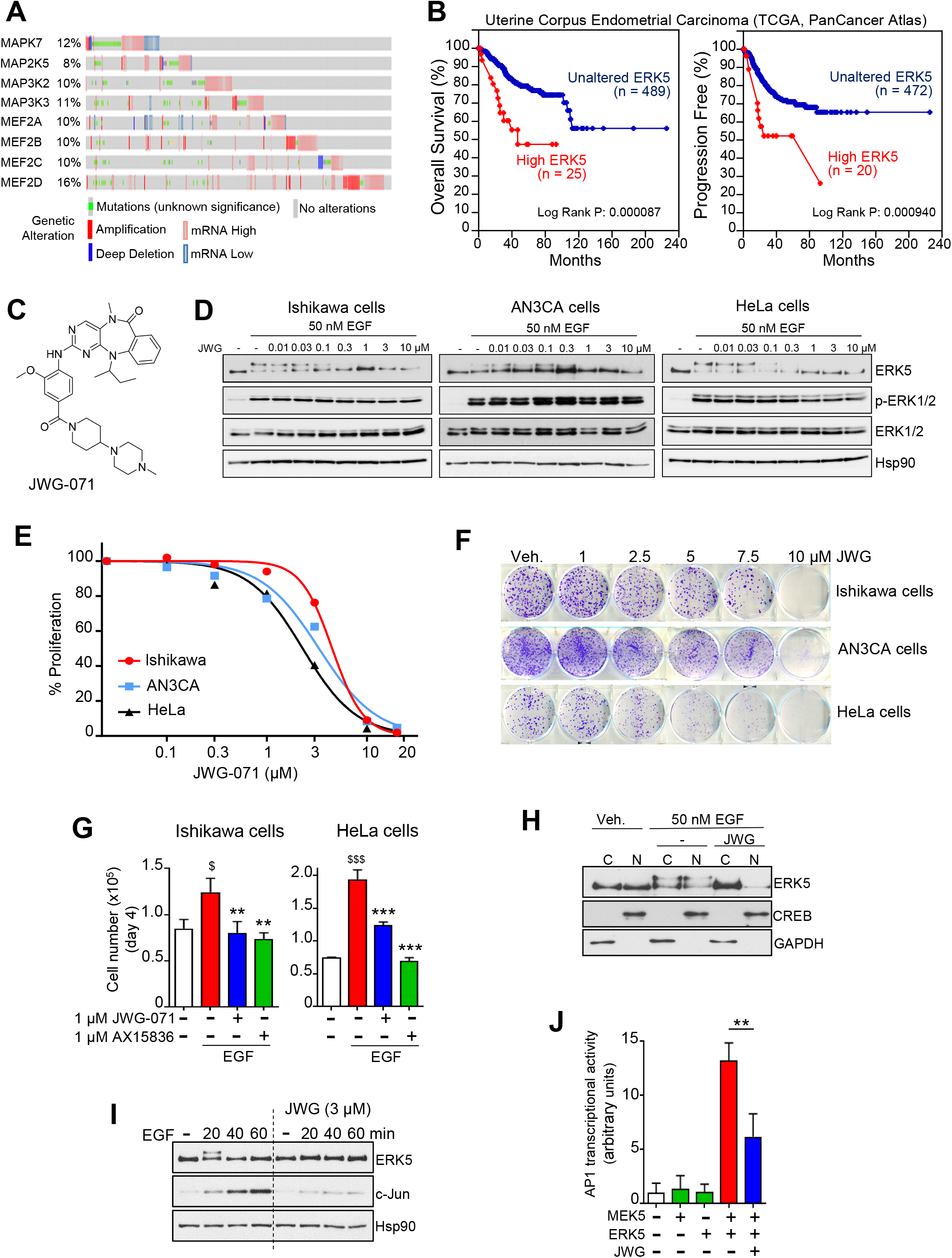
ERK5 inhibition impairs EC cell proliferation. **A**, Genomic profiles of components of the MEK5-ERK5 pathway in EC patients obtained from Uterine Corpus Endometrial Carcinoma data set (TCGA, PanCancer Atlas). Data set from cBiportal (https://www.cbioportal.org/). *MAPK7*, ERK5 gene; *MAP2K5*, MEK5. **B**, Kaplan-Meier plots overall survival (left) and progression free (right) in uterine corpus endometrial carcinoma patients with high (red), quartile 1) or normal (blue, quartile 2-3) ERK5 mRNA levels (data set from cBioportal). P-values were obtained using log-rank test. **C**, Chemical structure of JWG-071. **D**, Immunoblot analysis of the effect of JWG-071 on EGF stimulation. **E**, MTT cytotoxicity (48-hours) assay in a panel of human EC cells. **F**, 14-days clonogenic assay. **G**, Effect of ERK5 inhibitors JWG-071 and AX15836 in EGF-induced cell proliferation. Histograms show number of cells at day 4 referred to cells at day 0. ^$^, *P* < 0.05; ^$$$^, *P* < 0.001 EGF vs. vehicle. **, *P* < 0.01; ***, *P* < 0.001 ERK5i vs. EGF. **H**, JWG-071 blocks EGF-induced ERK5 nuclear localization. Cells were pre-incubated with 3 μM JWG-071 prior stimulation with 50 nM EGF (30 min), fractionated into the cytosolic and nuclear fractions, and protein expression analyzed by immunoblot. Hsp90, cytosolic marker; CREB-1, nuclear marker. **I**, ERK5 inhibition impairs EGF-induced c-Jun expression. Immunoblot analysis. **J**, JWG-071 impairs ERK5-mediated AP-1 transcriptional activity. Cells transfected with vectors encoding pAP-1–luciferase reporter and pRL-CMV-Renilla were subjected to a dual-luciferase reporter assay. **, *P* < 0.005.

### ERK5 inhibition impairs basal and EGF-induced EC cell proliferation

Early work reported that ERK5 mediates serum- and EGF-induced proliferation of cervical cancer HeLa cells [5], by phosphorylating and promoting transcriptional activation of MEF2C and transcription of c-*Jun* [16]. However, recent development of ERK5 inhibitors has led to question about the role of ERK5 kinase activity in cancer cell proliferation [17]. This is due to the discovery that widely used ERK5 ATP-competitive classical inhibitors (such as XMD8-92 and ERK5-IN-1) are unspecific, since they bind the acetyl-lysine domain of Bromodomain-Containing Protein-4 BRD4, a ubiquitous regulator of transcription and mediator of cancer cell proliferation. To address this question, we used the benzo[e]pyrimido-[5,4-b]diazepine-6(11H)-one core to developed a new ATP-competitive ERK5 inhibitor (JWG-071, **Fig. 1C**), with no BRD4 inhibitory activity [10]. JWG-071 shows selectivity for ERK5 (IC_50_ 90 nM, *in vitro* kinase assay, **Supplementary Fig. 2**) over BRD4 (IC_50_ 6 μM) [10]. In this work, we present the preclinical development of JWG-071.

Endometrial carcinoma cells express EGFR, and they represent a good model to investigate the role of EGF in cancer cell proliferation. Therefore, to study the role of ERK5 in basal and EGF-induced proliferation, we tested the JWG-071 compound in two different human endometrioid cancer cell lines that show EGFR expression: Ishikawa and AN3CA cells. We also used HeLa cells as a control, which also express EGFR. JWG-071 potently inhibited EGF-induced ERK5 phosphorylation in the three cancer cell lines tested (active ERK5 autophosphorylates, resulting in a slower migrating band), without affecting EGF-induced ERK1/2 phosphorylation (**Fig. 1D**). Proliferation assays showed that JWG-071 impaired cell proliferation (**Fig. 1E**) and colony-formation (**Fig. 1F**) in the cell lines tested.

Kinase selectivity of JWG-071 was previously determined by profiling the inhibitor against a panel of 468 human kinases, using an in vitro ATP-site competition binding assay. Apart of ERK5, JWG-071 also inhibited the activity of Doublecortin Like kinases DCLK1 and DCLK2 [10]. In order to investigate whether the antiproliferative effect of JWG-071 was due to DCLK1/2 off-target inhibition, we used the specific inhibitor DLCK1-IN-1 (which targets both DCLK kinases [18]). DLCK1-IN-1 had no effect on the proliferation of Ishikawa or AN3CA cells (**Supplementary Fig. 3**), suggesting that DCLK1/2 do not mediate the cytotoxicity induced by JWG-071 in EC cells.

Next, we investigated the role of ERK5 activity in EGF-induced cell proliferation. Cell counting experiments showed increased number of Ishikawa or Hela cells in response to EGF, which was significantly impaired by 1 μM JWG-071. Similar results were obtained using 1 μM AX15836 (**Fig. 1G**), another ERK5 specific inhibitor recently reported. Active ERK5 translocates to the nucleus, where it phosphorylates and activates the MEF2C transcription factor, which in turn promotes c-*Jun* expression and cell proliferation [16]. In our cell models, EGF treatment resulted in a rapid (30 min) accumulation of active/phosphorylated nuclear ERK5, whereas incubation with JWG-071 abolished nuclear active/phosphorylated ERK5 (**Fig. 1H**). Accordingly, the ERK5i impaired EGF-induced expression of c-Jun (**Fig. 1I**) and AP-1 transcriptional activity (**Fig. 1J**). These results suggest that ERK5 mediates EGF-induced proliferation by activating the MEF2C-c-Jun axis, at least in endometrial and cervical cancer cells that express EGFR.

### CRISPR/Cas9 MEK5 Knockout cells show impaired basal and EGF-induced cell proliferation

To ensure that the antiproliferative effect of JWG-071 in endometrial and cervical cancer cells expressing EGFR was due to inhibition of ERK5, we analyzed the effect of MEK5 depletion by generating stable CRISPR/Cas9 *MEK5* knockout Ishikawa and HeLa cell lines. MEK5-ERK5 constitute a unique signaling module, since MEK5 is the only upstream activating kinase for ERK5, and MEK5 has ERK5 as its only known substrate [19]. Therefore, MEK5^-/-^ cells lack ERK5 kinase activity and represent a useful tool to investigate the impact of ERK5 activity on cell physiology. As expected, and in contrast to wild type cells, MEK5^-/-^ Ishikawa or MEK5^-/-^ HeLa cells did not show ERK5 activation/phosphorylation in response to EGF (**Fig. 2A**). Importantly, genetic deletion of MEK5 did not affect phosphorylation of ERK1/2, Akt or S6 proteins in response to EGF (**Fig. 2A**) or insulin (**Supplementary Fig. 4A**), and neither affected phosphorylation of stress-associated MAP kinases p38 and JNKs in response to osmotic stress (**Supplementary Fig. 4B**). These results confirm that genetic deletion of *MEK5* does not affect the activity of other MAP kinases or the activity of the Akt-mTORC1 axis.

**Figure 2.**
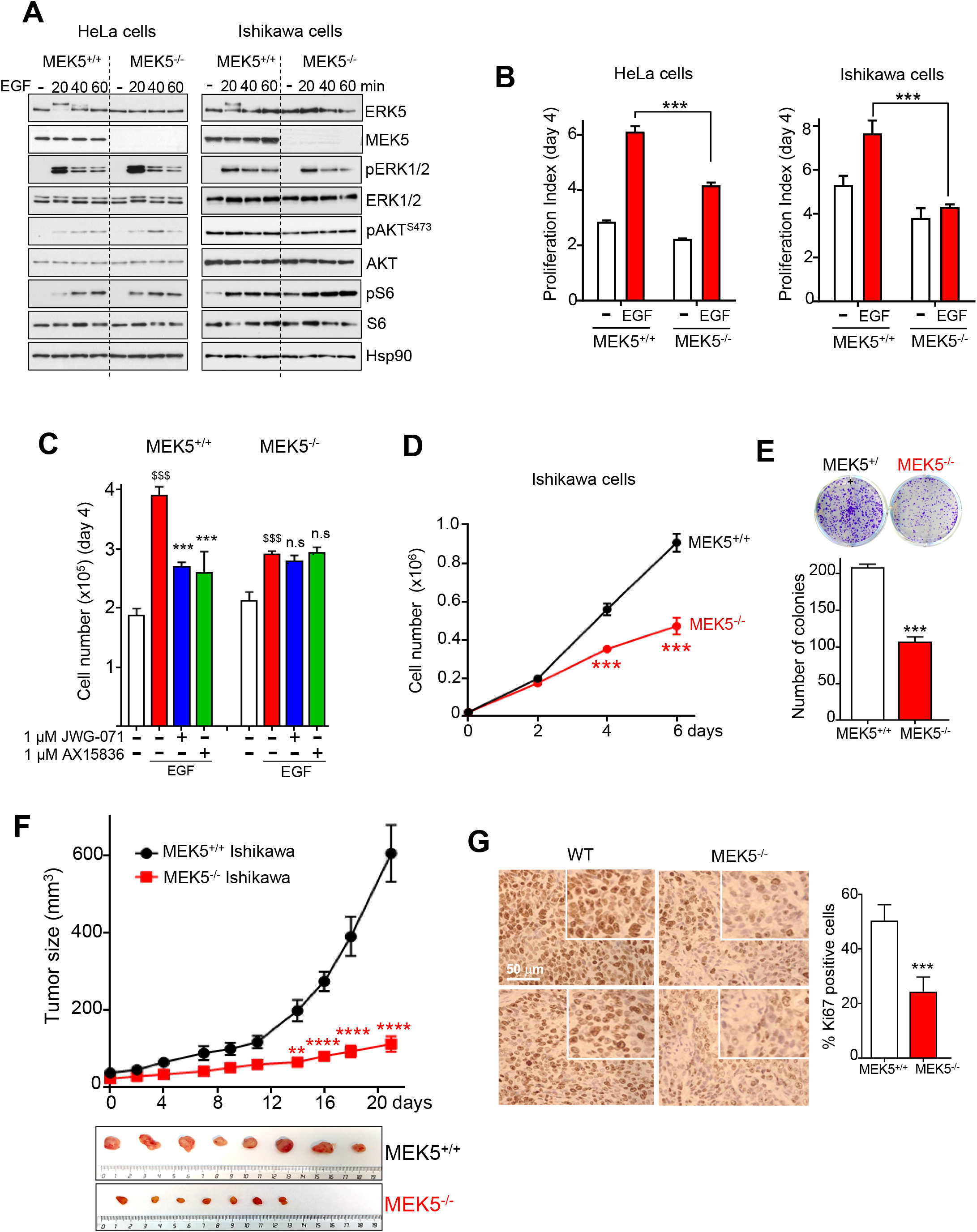
MEK5 genetic deletion impairs EC cell proliferation. **A**, CRISPR/Cas9 MEK5^-/-^ cells show impaired ERK5 activation in response to EGF stimulation. Cells were treated with 50 nM EGF and expression of proteins was monitored by immunoblot. **B**, CRISPR/Cas9 MEK5^-/-^ cells show impaired EGF-induced proliferation. Histogram shows the number of cells at day 4 referred to cells at day 0. **C**, ERK5 inhibitors do not impair EGF-induced proliferation in MEK5^-/-^ cells. ^$$$^, *P* < 0.001 EGF vs. vehicle. ***, *P* < 0.001 ERK5i vs. EGF. **D**, CRISPR/Cas9 MEK5^-/-^ Ishikawa cells show impaired proliferation, and impaired number of colonies. **E**, 14-day clonogenic assay. **F**, *In vivo* tumor growth of subcutaneous injected MEK5^+/+^ (n=8) and MEK5^-/-^ (n=7) Ishikawa cells. Data is presented as mean ± SEM. **, *P* < 0.01; ****, *P* < 0.0001, two-way ANOVA Bonferroni. **G**, Immunohistochemical analysis of Ki67 expression in MEK5^+/+^ and MEK5^-/-^ Ishikawa cell tumors. Right histogram shows the corresponding quantification. **B-D-E-G**, ***, *P* < 0.001.

As it happened for ERK5 inhibitor JWG-071, MEK5^-/-^ Ishikawa and MEK5^-/-^ HeLa cells showed significantly impaired proliferation in response to EGF in cell counting experiments (**Fig. 2B**). More importantly, concentrations of specific ERK5 inhibitors JWG-071 or AX15386 that impaired EGF-induced proliferation in wild type cells, had no effect in the absence of MEK5 (**Fig. 2C**). These results confirm that activation of the MEK5-ERK5 pathway is needed for EGF-induced proliferation. Regarding basal proliferation, MEK5^-/-^ Ishikawa cells phenocopied the effect of JWG-071, since they showed less proliferation (**Fig. 2D**) and colony-formation (**Fig. 2E**) capacity. Finally, to investigate the effect of *MEK5* deletion *in vivo*, wild type MEK5^+/+^ and MEK5^-/-^ Ishikawa cells were subcutaneously injected in athymic nude mice, and tumor growth was monitored. The depletion of MEK5 caused a drastic reduction on tumor growth *in vivo* (**Fig. 2F**), which was also evidenced by a reduction on the levels of the proliferation marker Ki67 (**Fig. 2G**).

In summary, our results support the use of JWG-071 as a specific ERK5 inhibitor for *in vitro* and *in vivo* studies. Furthermore, we provide evidence showing for the first time that ERK5 kinase activity mediates both, basal and EGF-induced proliferation in EC.

### ERK5 inhibition induces apoptosis in human EC cell lines

The above results prompted us to investigate the anticancer activity of ERK5 inhibition in EC. To do so, we used the endometrioid cancer cell lines Ishikawa and AN3CA cells (that carry PTEN deficiency and *P53* mutation), as well as the more clinically aggressive non-endometrioid serous adenocarcinoma cell line ARK1 (PI3KCA mutation). In cell viability assays, JWG-071 induced cytotoxicity in the three EC cell lines tested with IC_50_ values ranging 2-3 μM, as well as, in a wide panel of cancer cells, including prostate, cervical, neuroblastoma, and glioblastoma cancers (**Supplementary Fig. 5**).

Several authors have reported that the unspecific ERK5 inhibitor XMD8-92 induces apoptosis in different models of solid and blood cancers (reviewed in [9]). Therefore, we next analyzed whether JWG-071 induces apoptosis in EC cells. Flow cytometry analysis (Annexin V-PI staining) revealed that the ERK5i induced a significant increase of Annexin V staining in Ishikawa, AN3CA, and ARK1 cells (**Fig. 3A**). Furthermore, chromatin analysis revealed that JWG-071 treatment resulted in a drastic increase in number of cells with condensed and/or fragmented chromatin, a typical hallmark of apoptotic cell death (**Fig. 3B**). Immunoblot analysis confirmed that JWG-071 induced caspase-3 activation in the three EC cell lines tested, resulting in increased levels of cleaved/active caspase-3 and of cleaved PARP (caspase-3 canonical substrate) (**Fig. 3C**). To investigate the role of caspase activation in ERK5i-induced cancer cell death, we used the pan caspase inhibitor Q-VD-OPh. Preincubation with Q-VD-OPh significantly reverted the cytotoxicity of JWG-071 in Ishikawa, AN3CA, and ARK1 cells (**Fig. 3D**). We obtained similar results for the human cervical cancer cell line HeLa, where the ERK5i worked as a potent apoptotic inducer (**Supplementary Fig. 6**). Finally, we undertook a genetic approach to confirm that ERK5i induces apoptosis, by using apoptosis-deficient Bax/Bak double knockout MEF cells [20]. JWG-071 did not induce caspase activation (**Fig. 3E**) or cytotoxicity (**Fig. 3F**) in MEF Bax^-/-^/Bak^/-^, compared to MEF wild type cells. Thus, our results demonstrate that ERK5 inhibition induces apoptotic death in EC cells.

**Figure 3.**
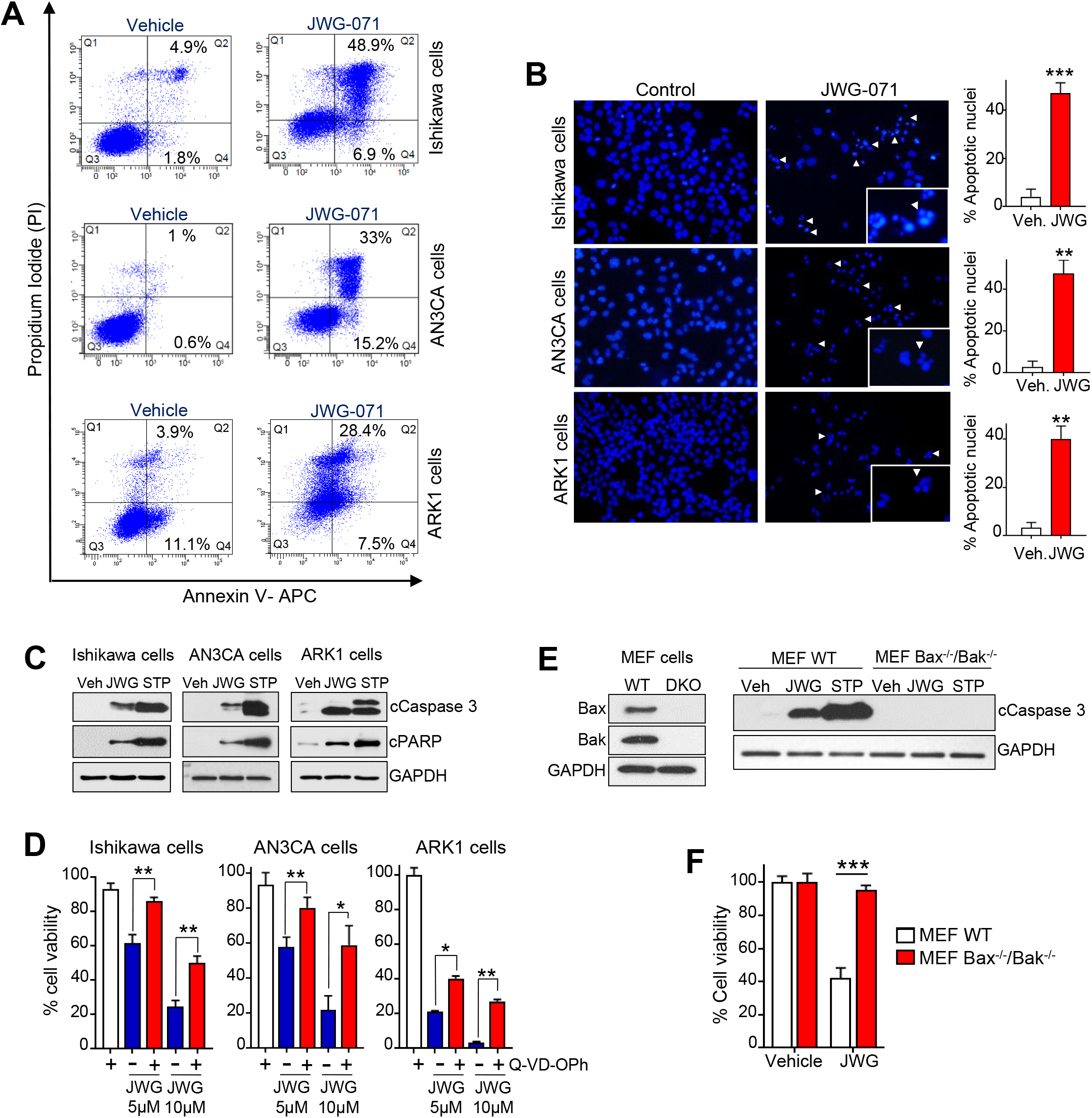
ERK5 inhibition induces apoptotic death in endometrioid and cervical cancer cells. **A**, Flow cytometry analysis of apoptosis induced by JWG-071 in EC cells. **B**, Representative images of nuclear morphology (Hoescht 33258) at 48-hours post-treatment with JWG-071 or vehicle. Arrowheads point condensed or fragmented nuclei. Right histograms, quantification of the results (four representative fields of each condition, n =3). **C**, Immunoblot analysis s of cells treated with vehicle or 5 μM JWG-071 (48 h). Staurosporine (STP, 18-hours) was used as a positive control. **D**, The pan-caspase inhibitor Q-VD-OPh impairs JWG-071-induced cytotoxicity. Cell viability (MTT) assay. **E-F**, Apoptosis-deficient cells are resistant to JWG-071 cytotoxicity. Bax^+/+^/Bak^+/+^ and Bax^-/-^/Bak^-/-^ MEF cells were treated with vehicle, 5 μM JWG-071 (48-hous) or 1 μM staurosporine (16-hours). Active caspase 3 (cCaspase 3) levels were monitored by immunoblot (**E**). Cell viability was determined by MTT assay (**F**). **B-D-F**, 0 *, *P* < 0.05; **, *P* < 0.005; ***, *P* < 0.001.

### ERK5 inhibition impairs NF-κB and activates JNK-apoptotic pathways in human endometrioid cells

Several authors have shown a role of the MEK5-ERK5 pathway in regulating the NF-κB canonical pathway in cancers such as leukaemia [21] and colon adenocarcinoma [22]. In these cancer models, ERK5 activity induces p65/RELA nuclear translocation and transcriptional activity, whereas ERK5 inhibition or silencing impairs p65/RELA activity. NF-κB pathway regulates, among other cancer hallmarks, proliferation and survival of numerous cancer types. Because it has been suggested that the NF-κB pathway controls proliferation and viability of EC cells [23], we next investigated the role of NF-κB pathway in the anticancer activity of ERK5i.

Gene expression data mining from the cBioPortal for Cancer Genomics (Uterine corpus endometrial carcinoma TCGA, PanCancer Atlas), using the database GEPIA2 (http://gepia2.cancer-pku.cn/#index), showed a strong positive correlation between *MAPK7* and *RELA* mRNA expression levels (R, 0.47; Spearman, 8.2e^−11^: **Fig. 4A**). We next analyzed ERK5 and p65/RELA protein levels in tumoral and non-tumoral (peritumoral) samples from 17 endometrioid cancer patients (**Supplementary Fig. 7A**). Immunoblot analysis revealed that 12 of 17 tumor samples showed increased p65/RELA (p:0.031) and ERK5 (p:0.0562) levels, compared with peritumoral tissue (**Fig. 4B and Supplementary Fig. 7B)**. No differences in ERK5 protein levels were observed among different cancer grades (**Supplementary Fig. 7C**). Of note, we found a strong correlation between relative endometrial tumoral/non-tumoral ERK5 and p65 levels (R:0.835, p:0.00003, **Fig. 4C**).

**Figure 4.**
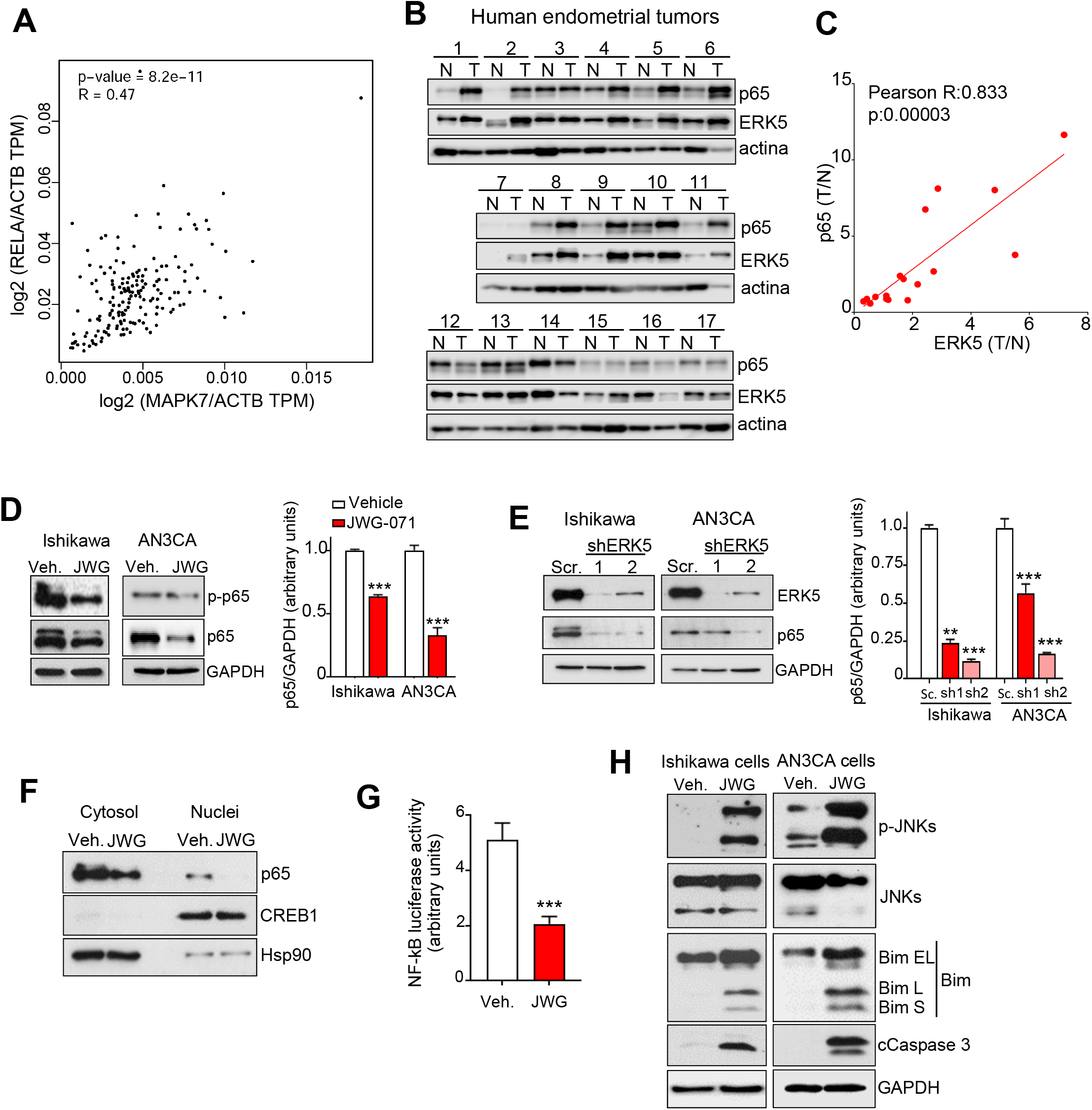
ERK5 inhibition impairs NF-kB pathway and JNK-induced apoptosis in endometrioid cancer cells. **A**, Spearman correlation analysis of ERK5 and p65/RELA mRNA expression in uterine corpus endometrial cancer patients was analyzed and plotted by GEPIA website (http://gepia.cancer-pku.cn/index.htlm). **B**, Immunoblot analysis of samples from 17 different patients with endometrioid cancer. T, tumor area; N, non-tumor area. **C**, Pearson’s pairwise correlation analysis of relative levels of ERK5 and p65 protein expression in non-tumoral and tumoral endometrial samples. **D**, ERK5 inhibition impairs p65 levels in EC cells. Right histograms show the quantification of the results. **E**, ERK5 silencing impairs endogenous p65 levels. Immunoblot analysis. Cells were infected with lentiviral shRNA particles encoding for either scramble or two different shERK5. Right histogram shows the quantification of p65 expression. **F**, ERK5 inhibition impairs nuclear p65. Ishikawa cells treated with JWG-071 for were fractionated to obtain cytosolic and nuclear fractions. Expression of indicated proteins was monitored by immunoblot. **G**, ERK5 inhibition impairs NF-κB transcriptional activity. Cells were transfected with plasmids encoding for a Luciferase-NF-κB reporter and pRL-CMV-Renilla, treated with JWG-071 for 24 h, and subjected to dual-luciferase reporter assay. **H**, ERK5 inhibition activates JNK apoptotic pathway. Cells were treated with vehicle or JWG-071 (48 h) and proteins analyzed by immunoblot. **D-E-G**, **, *P* < 0.005; ***, *P* < 0.001.

Next, we investigated the impact of ERK5 inhibition on the NF-κB canonical pathway in EC cells. In Ishikawa and AN3CA EC cells, the ERK5i induced a significant reduction of p65 and phospho-p65 levels (**Fig. 4D**), whereas ERK5 silencing with two different shRNAs also induced a significant decrease in p65 protein levels (**Fig. 4E**). Moreover, the ERK5i treatment reduced nuclear p65 levels (subcellular fractionation assay, **Fig. 4F**) and NF-κB transcriptional activity (**Fig. 4G**). Since impairment of NF-κB canonical pathway results in activation of the JNK-mediated apoptotic pathway in several cancers (21-22), we investigated if this was also the case in our EC cell models. Treatment of Ishikawa and AN3CA cells with the ERK5i resulted in activation/phosphorylation of JNKs, and in the appearance of Bim-L and Bim-S, the more cytotoxic forms of the pro-apoptotic JNK substrate (**Fig. 4H**).

To explore the role of the NF-κB pathway in endometrioid cancer cell viability, we investigated the effect of inhibition of NF-κB. The IκB kinase (IKK) inhibitor BAY11-7082 induced cytotoxicity (MTT assay) and apoptosis (flow cytometry assay) in Ishikawa and AN3CA cells, with IC_50_ values ranging 2-3 μM (**Fig. 5A-B**). Furthermore, as it happened for JWG-071, treatment of EC cell lines with BAY11-7082 resulted in impaired p65 phosphorylation and activation of the JNK proapoptotic pathway (**Fig. 5C**). Parallel experiments demonstrated that overexpression of the dominant negative mutant SR-IκBα (a non-degradable form of IκBα, [13]) impaired p65 expression and activated the JNK-apoptotic pathway (**Fig. 5D**). Of note, these results also suggest an important role of the NF-κB pathway in EC cell viability.

**Figure 5.**
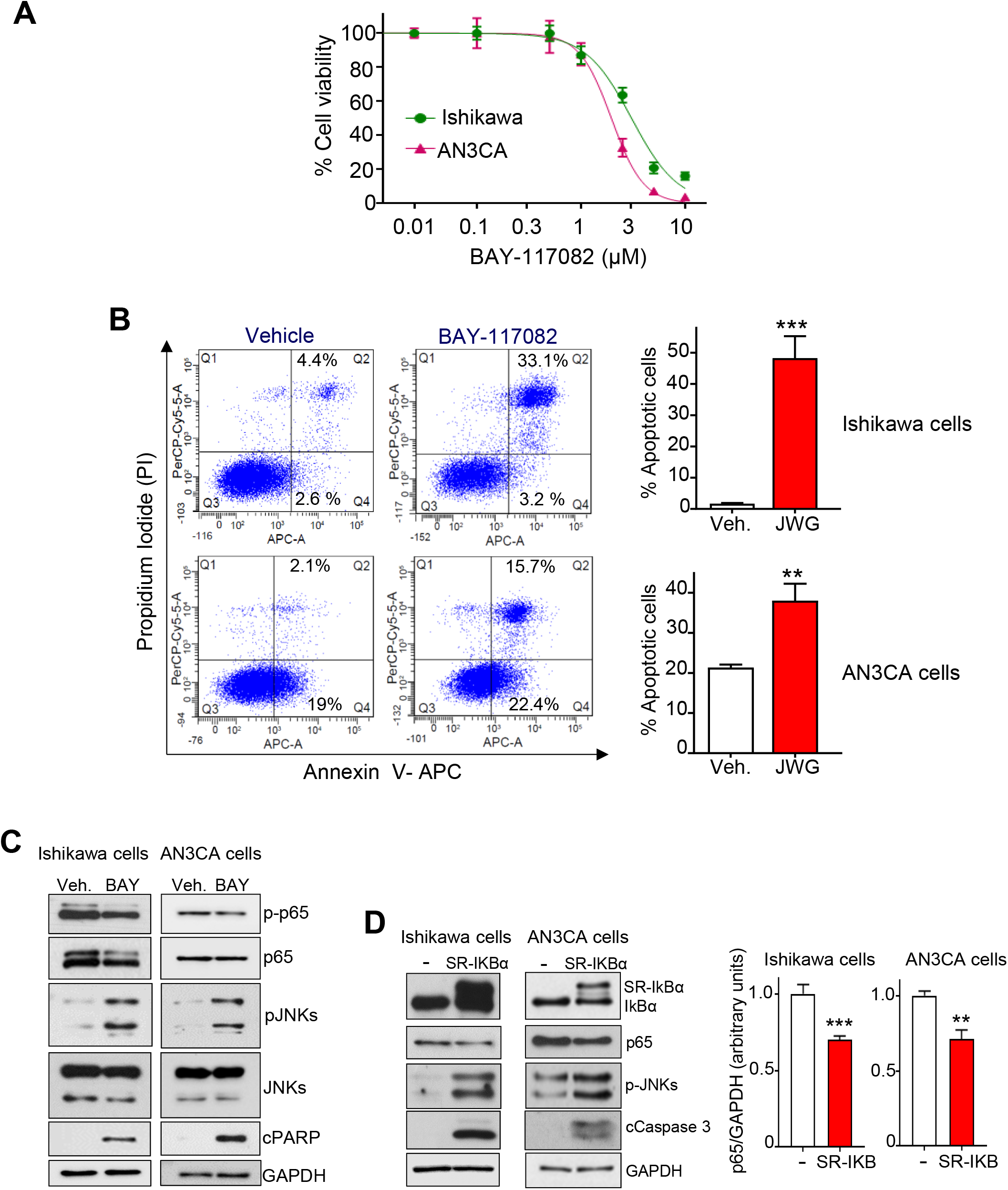
The NF-kB inhibition induces apoptotic cell death in endometrioid cancer cells. **A**, Ishikawa and AN3CA cells were treated with BAY11-7082 (NF-kB pathway inhibitor) for 48 h and cell viability was assessed by MTT assay. Similar results were obtained in three separate experiments. **B**, Representative results of flow cytometry analysis of BAY11-7082-induced cell apoptosis. Ishikawa or AN3CA cells treated with BAY11-7082 10 µM for 48h were stained with Annexin V and propidium iodide (left panel). Histograms represent the percentage of apoptotic cells (early and late apoptosis), obtained in response to vehicle or BAY11-7082 treatment. Mean values ± SD of assays performed in duplicate are shown. **C**, Expression of proteins from cells treated with NF-kB inhibitor BAY-118072 (10 μM, 48 h) was evaluated by immunoblotting. **D**, Impairment of p65/RELA expression activates JNK apoptotic pathway. Immunoblot analysis of cells transiently transfected with an empty vector or a vector encoding for a non-degradable IKBα protein (SR-IKBα). Right histograms show the corresponding quantification of p65 levels, referred to GAPDH. *, *P* < 0.05; **, *P* < 0.01; ***, *P* < 0.001 (Student’s t-test).

### IKKɣ/NEMO mediates ERK5i-induced EC cell death

To investigate the mechanism involved in ERK5i-induced p65/RELA impairment, we next analyzed the expression levels of proteins of the NF-κB canonical pathway. In response to cytokines such as TNFα or IL-1, the NF-κB essential modulator NEMO/IKKγ recruits and allows the activation of the IkB kinases IKKα and IKKβ Phosphorylated IkB undergoes ubiquitylation and proteasomal degradation, allowing nuclear translocation of the NF-kB complex (p50/p65) and activation of transcription [26]. ERK5i treatment resulted in a drastic reduction of NEMO/IKKγ protein levels, as well as in reduced IKKα/IKKβ/IkBα levels (**Fig. 6A**). ERK5i also impaired *NEMO/IKKγ* mRNA levels in EC cells, suggesting a role for ERK5 in *NEMO*/*IKKγ* transcription (**Fig. 6B**).

**Figure 6.**
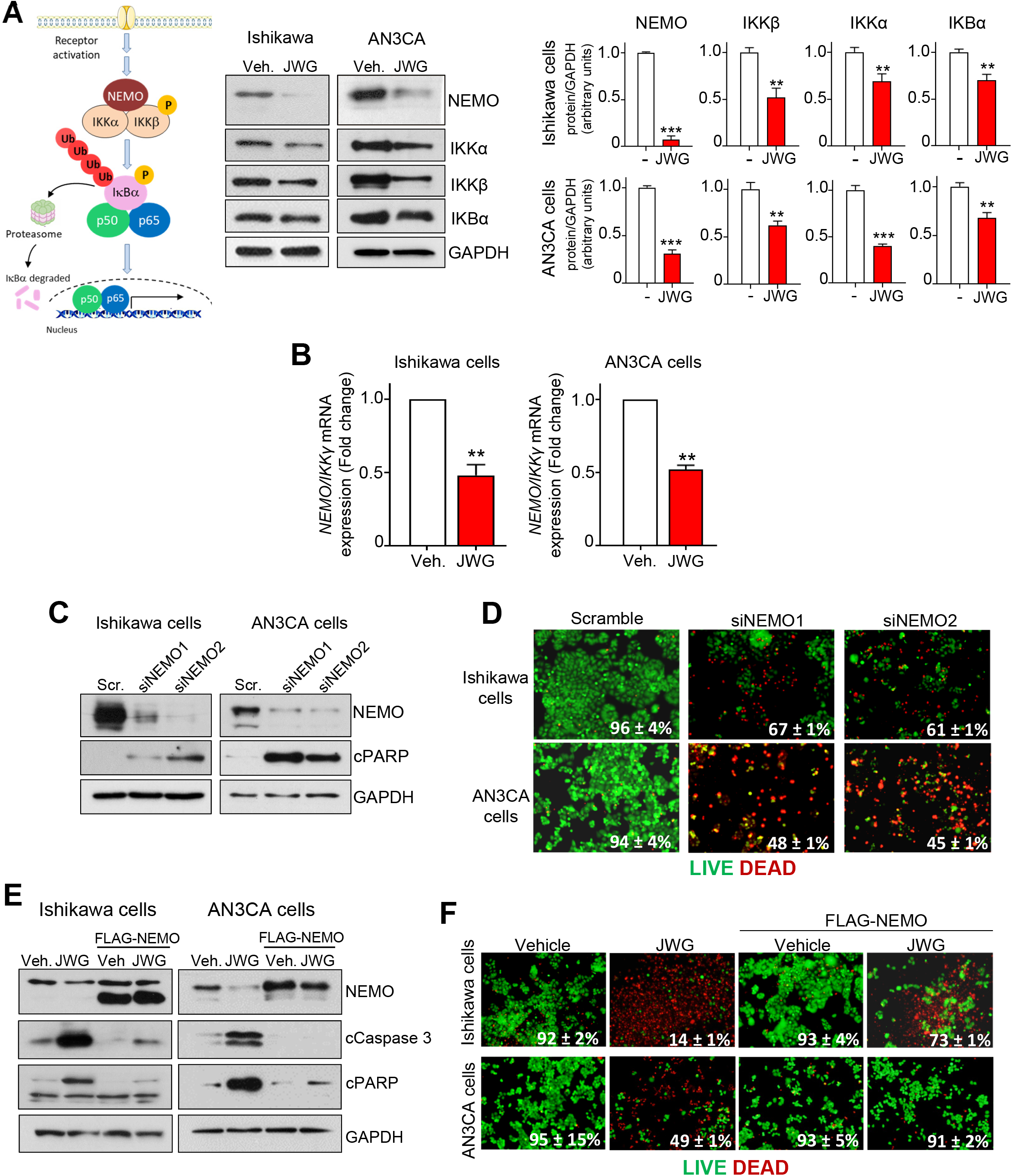
IKKɣ/NEMO mediates ERK5i-induced EC cell death. **A**, ERK5 inhibition decreases protein levels of IKKɣ/NEMO and downstream components of the NF-kB canonical pathway. Protein levels of cells treated with either vehicle or JWG-071 (48h) were analyzed by immunoblotting. Histograms show the quantification of the results. **B**, ERK5 inhibition impairs NEMO/IKKɣ mRNA levels. mRNA levels were analyzed by qRT-PCR and normalized by GAPDH mRNA levels. **C**, NEMO/IKKγ silencing induces apoptosis. EC cells were transfected with control siRNA (Scr) or *NEMO/IKK*γ-selective siRNAs and levels of proteins were monitored by immunoblotting. **D**, *NEMO/IKKγ* silencing induces EC cell death. Cells were stained with LIVE/DEAD reagent, and alive (green) and dead (red) were visualized by fluorescence microscopy. Figure shows representative fields, and percentage of alive cells is given. **E**, NEMO/IKKγ overexpression prevents caspase 3 activation in response to JWG-071. Cells transfected with either an empty vector or a vector encoding for human NEMO/IKKγ were treated with vehicle or JWG-071 (48 h), and levels of the indicated proteins were monitored by immunoblotting. **F**, NEMO/IKKγ overexpression prevents EC cell death induced by JWG-071. **A-B**, **, *P* < 0.005; ***, *P* < 0.001.

Next, we studied the impact of NEMO in EC cell viability. Silencing of *NEMO/IKKγ* with two different siRNAs resulted in activation of caspase-3, as judged by the cleavage of the caspase-3 substrate PARP (**Fig. 6C**) and in a drastic reduction of cell viability (**Fig. 6D**). Conversely, overexpression of NEMO protein protected EC cells from the cytotoxicity and apoptosis (active caspase-3) induced by the ERK5i (**Fig. 6E-F**). Of note, DLCK1 inhibition did not affect NEMO/IKKγ or p65 protein expression levels in Ishikawa or AN3CA cells (**Supplementary Fig. 8**). All in all, these results suggest that NEMO/IKKγ expression mediates the cytotoxicity induced by ERK5 inhibition in EC cells.

### Antitumor activity of ERK5i in EC xenografts

To determine the antitumor activity of JWG-071 *in vivo*, we selected the Ishikawa cells as a representative EC cell line. The suitability of JWG-071 for *in vivo* use was assured by pharmacokinetic analysis. JWG-071 demonstrated a favourable pharmacokinetic profile in mice, with 84% oral bioavailability, a T_max_ of 1 h, and a half-life of 4,34 h (**Supplementary Fig. 9**). We first tested the effect of JWG-071 systemic administration on the proliferation of EC tumors in short-term experiments. JWG-071 significantly impaired the growth of human endometrial tumor xenografts (Ishikawa cells) after 5- or 7-days treatment (**Fig. 7A**). Immunohistochemical analysis showed a significant decrease in levels of the proliferation marker Ki67 in tumors treated with the ERK5i, suggesting that JWG-071 treatment impaired the proliferation of EC tumor cells *in vivo* (**Fig. 7B**).

**Figure 7.**
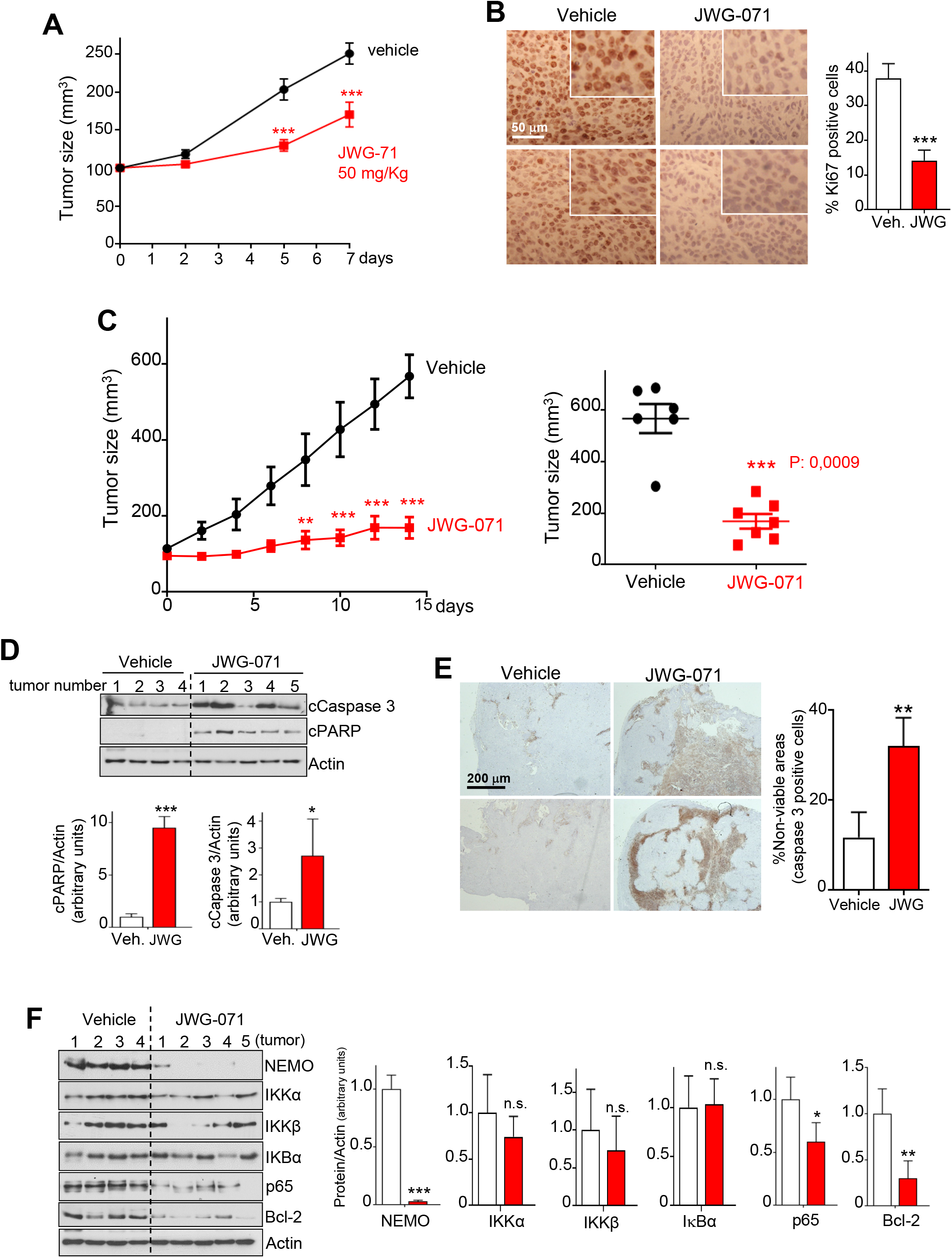
ERK5 inhibition impairs growth of EC tumor xenografts *in vivo*. **A**, Tumor growth curve of subcutaneous xenograft (Ishikawa cells), treated i.p. with either vehicle or 50 mg/Kg JWG-071 daily. N=6 mice per group. Data is presented as mean ± SEM. ****, *P* < 0.0001, two-way ANOVA Bonferroni. **B**, Immunohistochemical analysis of Ki67 expression of xenograft tumors. Right histograms show Ki67 quantification. ***, *P* < 0.001. **C**, Tumor growth curves. Nude mice bearing Ishikawa cell subcutaneous xenograft tumors were daily treated i.p. with either vehicle (n=6) or 50 mg/Kg JWG-071 (n=6). Values represent the mean ± SEM. *, *P* < 0.05; ***, *P* < 0.001, two-way ANOVA Bonferroni. Right panel shows the corresponding scatter plot for tumors at day 14 of treatment. **D-E**, Immunoblot **(D)** and cleaved caspase 3 immunohistochemical **(E)** analysis of the endometrioid xenograft tumors. Histograms show the corresponding quantification. **F**, Immunoblot analysis of NF-κB pathway proteins from xenograft tumors. Histograms show the corresponding quantification. **B-D-E-F**, *, *P* < 0.05; **, *P* < 0.01; ***, *P* < 0.001.

In long-term experiments in mice with established subcutaneous Ishikawa tumors, JWG-071 (50 mg/kg once per day) potently impaired tumor growth, with ∼85% tumor growth inhibition (**Fig. 7C**), without affecting mouse weight (**Supplementary Fig. 10**). Immunoblot (**Fig. 7D**) and immunohistochemical (**Fig. 7E**) analysis showed that administration of JWG-071 potently activated apoptosis in EC xenograft tumors. Furthermore, ERK5i treatment also resulted in a drastic impairment of NEMO/IKKγ and p65/RELA protein levels in tumors, as well as of the p65/RELA transcriptional substrate Bcl-2 (**Fig. 7F**). No significant changes in IKKα/IKKβ/IkBα protein levels were observed. Together, these results demonstrate that effective ERK5 inhibition impairs the NF-κB canonical pathway, as well as cell and tumor viability.

### ERK5i sensitizes endometrioid cancer cells and tumor xenografts to chemotherapy

Recent evidence has led to propose ERK5 inhibition as a promising strategy to sensitize lung, breast, or colorectal cancer cells to standard chemotherapy (see [8] for review). To investigate whether this was also the case for EC, we used paclitaxel and carboplatin, two chemotherapics used in combination as standard of care to treat patients with advanced EC (1). ERK5 inhibition synergized with either paclitaxel or carboplatin to promote cytotoxicity in Ishikawa and AN3CA EC cells (**Fig. 8A**, see combination index values lower than 1). We obtained similar results in colony-formation assays, where combined ERK5i and chemotherapy treatment resulted in a drastic reduction in the number of colonies, compared to single treatments (**Fig. 8B**). Importantly, CRISPR MEK5^-^/^-^ cells, which lack ERK5 kinase activity, also showed enhanced sensitivity to paclitaxel and carboplatin treatment, reinforcing the notion that pharmacologic (ERK5i) or genetic (MEK5 knockout) inhibition of ERK5 kinase activity sensitizes EC cells to standard chemotherapy (**Fig. 8C**). Of note, the NF-κB inhibitor BAY11-7082 also sensitized Ishikawa and AN3CA EC cells to paclitaxel and carboplatin (**Supplementary Fig. 11**).

**Figure 8.**
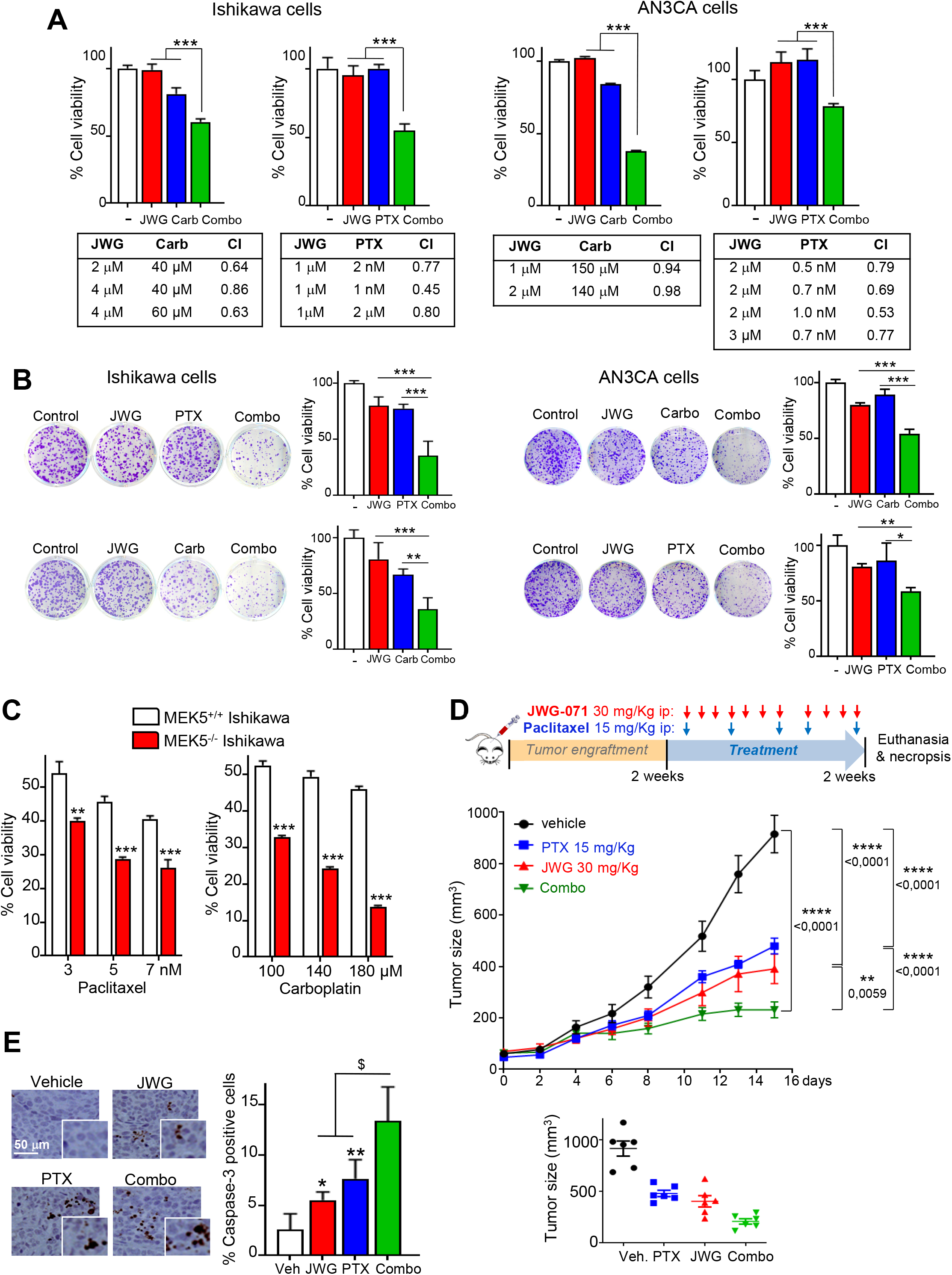
ERK5 inhibition sensitizes endometrioid cancer cells to standard chemotherapeutic agents. **A**, Cells were treated with the indicated concentrations of JWG-071, cisplatin (CPT), carboplatin (Carb) or paclitaxel (PTX) for 48 h, and cell viability was determined by MTT assay. Lower tables show the combination index (CI) analysis, obtained using the Compusyn software (CI > 1, antagonism; CI = 1, summary effect; CI < 1, synergism). **B**, 14-day clonogenic assay. Left, representative images of single-cell clone proliferation, stained with crystal violet. Right, quantification of number of colonies. **C**, *MEK5* deletion sensitizes EC cells to chemotherapy. Cell viability (MTT) assay of *MEK5*^+/+^ and *MEK5*^-/-^ Ishikawa cells. **A-B-C**, *, *P* < 0.05; **, *P* < 0.005; ***, *P* < 0.001. **D**, ERK5i sensitizes EC tumor xenografts to chemotherapy. Athymic nude mice bearing EC tumor xenografts (Ishikawa cells) were treated with vehicle, 30 mg/Kg JWG-071 (i.p., daily), 15 mg/Kg paclitaxel (i.p., BIW) or combo. Values represent the mean ± SEM of 6 mice for each group. Low panel shows the corresponding scatter plot for tumors at day 15 of treatment. **, *P* < 0.05; ****, 0.0001, two-way ANOVA Tukey. **E**, Tumors were collected and cleaved caspase 3 was evaluated by immunohistochemical analysis. Right histogram shows the corresponding quantification. ^$^, *P* < 0.05 combo treatment vs. single treatments. * *P* < 0.05; ** *P* < 0.001 ERK5i or paclitaxel vs. vehicle.

Finally, to evaluate the effectiveness of ERK5 inhibition and chemotherapy as a combinatorial treatment *in vivo*, we investigated the effect of JWG-071 and/or paclitaxel in athymic nude mice subcutaneously injected with human endometrioid Ishikawa cells. Single treatments with low doses of paclitaxel (15 mg/Kg) or of JWG-071 (30 mg/Kg) produced 40-50% tumor growth inhibition, compared to the control group. However, combinatorial treatment using both paclitaxel and JWG-071 resulted in a significantly greater inhibition of tumor growth compared with vehicle-treated, tumor-bearing mice (combo vs paclitaxel p<0.0001; combo vs JWG-071 p=0.0059) (**Fig. 8D**). Both mono and combinatorial treatments were well tolerated by mice, without significant weight loss or other overt effects (**Supplementary Fig. 10B**). In line with results obtained in EC cultured cells, tumors from mice receiving combinatorial therapy exhibited significantly greater staining for the apoptotic marker active caspase-3, compared with tumors from mice receiving single treatment (**Fig. 8E**). Thus, ERK5 inhibition sensitizes EC cells and tumors to standard chemotherapy *in vitro* and *in vivo*.

## Discussion

EC is the most common gynaecological malignancy in the western world, accounting for one third of the deaths related to gynaecological cancers [1]. Early-stage EC carcinomas are treated by surgery and radiation, but the advanced stage has a very poor prognosis (1). Standard pharmacological treatment of advanced endometrial carcinoma uses a combined regimen of taxanes (paclitaxel) and platinums (carboplatin) (1). Often, treatments fail due to the emergence of chemoresistance, and other therapeutic options are limited. Therefore, there is an urgent need for new therapeutic therapies to improve the treatment of advanced EC. Understanding the signal transduction pathways dysregulated in EC might be of help in identifying novel therapeutic targets to improve patients’ prognosis and treatment outcomes.

In this study, we propose ERK5 inhibition as a new potential targeted therapy to treat EC, both as monotherapy and in combination with standard chemotherapy. Hence, we report the preclinical development of the small-molecule JWG-071, a potent and selective ERK5 inhibitor with a good pharmacokinetic profile. JWG-071 is the first specific ERK5 kinase inhibitor (no BRD4 activity) that shows antitumor activity in animal models. Specifically, we demonstrate the requirement of ERK5 kinase activity for EC cell proliferation and survival. In line with this, EC patients with high levels of ERK5 mRNA expression had significant lower overall survival and progression-free survival (Uterine corpus endometrial carcinoma TCGA, PanCancer Atlas, cBioPortal).

Early work reported a role for ERK5 in mediating EGF- and serum-induced proliferation in cervical cancer cells [5]. Interestingly, EGFR is commonly expressed in normal endometrium, and its overexpression in EC (43-67% of EC patients) is associated with advanced stage and poor prognosis [27]. Here, we show for the first time that ERK5 inhibition impaired basal and EGF-induced proliferation of EC cells that express EGFR (**Fig. 1-2**). Specific ERK5 inhibition was achieved using either the JWG-071 small compound or EC cells lacking the ERK5 upstream kinase MEK5. MEK5^-/-^ EC cells showed a reduced basal proliferation activity and clonogenicity, as well as a drastic reduced tumor growth in nude mice (**Fig. 2**), phenocopying the effect of the ERK5i. Importantly, MEK5 deletion in EC cells did not affect activation of ERK1/2, Akt or mTORC1 in response to EGF or insulin, nor did it affect basal activity of these kinases. Our results suggest a genuine role of the MEK5-ERK5 pathway in mediating EC proliferation, at least in epithelial EC cells that express EGFR, as it has been reported for other carcinoma cancers such as hepatocellular carcinoma [28], prostate adenocarcinoma [29] or non-small cell lung cancer [30]. Of note, it has been reported that ERK5 is dispensable for proliferation of colorectal cancer cells carrying *KRAS/BRAF* mutations [31], suggesting that genetic mutational background must be considered in order to establish the exact role of ERK5 in proliferation of different cancer types.

Mechanistically, ERK5 inhibition of EC cells resulted in impairment of EGF-induced c-Jun expression and AP-1 transcriptional activity, as reported for other ERK5i in cancer types such as breast cancer [32] and hepatocellular carcinoma [28]. Thus, ERK5 inhibition, by preventing EGF-induced ERK5 nuclear translocation, impaired the ERK5-MEF2C-c-Jun axis, leading to slowing down proliferation.

ERK5 also participates in the proliferative signaling in response to activation of HER2 receptors in breast cancer cells, [33], and ERK5 inhibition sensitizes these cells to anti-HER2 treatments [34]. 20%-40% of patients with advanced non-endometrioid EC show HER2 overexpression [35], and Trastuzumab added to standard chemotherapy improved survival of patients with HER^+^ aggressive serous EC, according to a recent phase 2 study [36]. Thus, it will be relevant to investigate whether ERK5 also mediates HER2 signaling in aggressive EC, and whether ERK5 inhibition could be of help to sensitize these cells to anti-HER2 therapies.

The NF-κB pathway is a key regulator of proliferation and survival of many types of cancers, by controlling the expression of a plethora of genes involved in proliferation, angiogenesis, inflammation, and survival, among others [37]. Early work suggested a role for the NF-κB pathway in endometrial carcinogenesis [23]. This was later confirmed by an IHC study that found increased nuclear NF-κB expression in patients with EC, compared with benign and hyperplasic endometrial tissues, and suggested a role for NF-κB pathway in the development of malign endometrial changes [38]. These observations are in agreement with those reported for other gynaecological cancers, such ovary [39] and cervical [40] cancers, where NF-κB is involved in tumor progression.

Here, we found a significant and positive correlation between ERK5 and p65/RELA protein levels in human EC tumor samples, and that ERK5 inhibition or silencing resulted in impairment of the NF-κB canonical pathway in EC cells and tumor xenografts. These results are in agreement with those reported for colorectal [22] and leukemic [21] cancers, where MEK5/ERK5 overexpression results in enhanced p65/RELA nuclear localization and transcriptional activity. Furthermore, ERK5 is essential for NF-κB-induced survival in leukemic cells [21], and ERK5 protein levels positively correlate with p65/RELA levels in colon carcinoma tumor samples [22]. However, the mechanism by which ERK5 modulates NF-κB pathway in several cancer types remains to be described. In this work, we report that ERK5 inhibition or silencing results in downregulation of NEMO/IKKγ expression, leading to impaired p65/RELA activity and apoptosis, which was rescued when NEMO/IKKγ was transiently overexpressed. Although we found that ERK5 inhibition significantly reduced NEMO/IKKγ mRNA levels, we cannot discard a role of ERK5 in regulating NEMO/IKKγ stability, as described for other ERK5 substrates such as KLF2 [41]. Further work will be necessary to establish if ERK5 phosphorylates NEMO/IKKγ, and the implications in the stability of this protein.

We also provide evidence showing, for the first time, that the NEMO-IKKβ-p65 axis mediates EC cell survival. Hence, NEMO silencing, IKKβ inhibition, or p65 inhibition (by the SR-IkB repressor) resulted in apoptotic death of EC cells. Our results point to NF-κB as an interesting target for EC treatment, as it has been proposed for other cancer types [37]. However, several IKKα or IKKβ inhibitors have failed in clinical trials, mainly due to toxicity concerns [42]. This was probably due to the fact that these inhibitors block the activation of the IKK complex (IKKα/IKKβ, necessary for the activation of the canonical NF-κB pathway) as well as the activity of individual IKKβ and IKKα (necessary for the activation of the non-canonical pathway) [42]. On the contrary, NEMO/IKKγ inhibition affects the activity of the NF-κB pathway mediated by the IKK complex (canonical pathway), but not the activity of individual IKKα or IKKβ or the non-canonical pathway. Concordantly, a small compound that blocks the binding of NEMO/IKKγ to IKKβ has shown anticancer activity in colorectal cancer tumor xenografts [43]. Considering that ERK5 inhibition or silencing impaired NEMO/IKKγ expression in EC cells and xenograft tumors, we speculate that ERK5 inhibition could be an effective and safe strategy to impair the NF-κB pathway in those cancers where NF-κB acts as a driver, such as EC.

Finally, here we report that ERK5 inhibition sensitizes EC cells and xenograft tumors to the standard chemotherapy agents paclitaxel and carboplatin (1), suggesting that ERK5i could be used as an effective targeted therapy to improve chemotherapy of EC patients. Our results are in line with those reported for other cancer types. Thus, among others, ERK5 inhibitor XMD8-92 improves the cytotoxicity of docetaxel and/or doxorubicin in lung [44], triple negative breast [45] and squamous skin [46] cancer cells and tumors, as well as the antitumor action of 5’-fluoroacil in p53-dependent colorectal cancer cells [47]. Furthermore, ERK5 silencing also potentiates doxorubicin and cisplatin therapies in malignant mesothelioma cells [48], or doxorubicin, cyclophosphamide, and paclitaxel in TNBC cells [49]. The mechanism by which ERK5 inhibition sensitizes several types of cancer cells remains to be described, although it has been suggested that it could be due to inhibition of some of the ABC transporters [50]. Since we found that pharmacologic inhibition of the NF-κB pathway (IKKβ inhibitor BAY-117082) also potentiated the cytotoxic effect of paclitaxel and carboplatin in EC cells, it is plausible that NF-κB inhibition may mediate the sensitization of EC cells to chemotherapy exerted by ERK5 inhibitors. Moreover, because activation of the NF-κB pathway is implicated in resistance to taxanes and platinum-based drugs in several preclinical models of cancer [51], it is tempting to speculate that ERK5 inhibition might escape from NF-κB-mediated resistance to chemotherapy in EC cells. Further work will be necessary to investigate this possibility.

## Acknowledgments

S. Espinosa-Gil is a recipient of a fellowship from FI-AGAUR (2020-FISDU-00575). N. Dieguez-Martinez and I. Bolinaga-Ayala are recipients of PIF fellowships from UAB. We are grateful to Cristina Gutierrez, and Neus Ontiveros for tissue culture, to Mar Hernández for IHC assistance. We also thank Miguel Segura and Victor Yuste for helpful discussions. We thank the following UAB Services: Servei de Cultius Cel·lulars INc, Laboratori de Luminiscència Espectroscopia and Servei Genòmica Informàtica. We also thank to the UAT Service from VHIR.

## Statements & Declarations

### Author Contributions

Conceived and designed the experiments: ND-M, SE-G, XD and JML. Performed experiments: : ND-M, SE-G, GY, EM-R, IB-A, MV-C, ID-O and JML. Analyzed the data: ND-M, SE-G, HP-M, EC, JRB, XD and JML. Writing–original draft: ND-M, SE-G and JML. All the authors read, critically revised, and approved the final manuscript

### Data Availability

The original contributions presented in the study are included in the article/supplementary material, and further inquiries can be directed to the corresponding author.

### Ethics approval

The animal study was approved by the Ethics Committee on Animal Experiments of the Universitat Autònoma de Barcelona (CEEAH) and the Ethics Commission in Animal Experimentation of the Generalitat de Catalunya, Spain.

### Consent to participate

The use of samples from patients was approved by the Institutional Review Boards of Hospital Vall d’Hebron (Barcelona, Spain), and written informed consent was obtained from the patients involved.

### Consent to publish

Not applicable

