## Supplemental Figures and Table for "The ERK5/NF-κB signaling pathway targets endometrial cancer proliferation and survival"

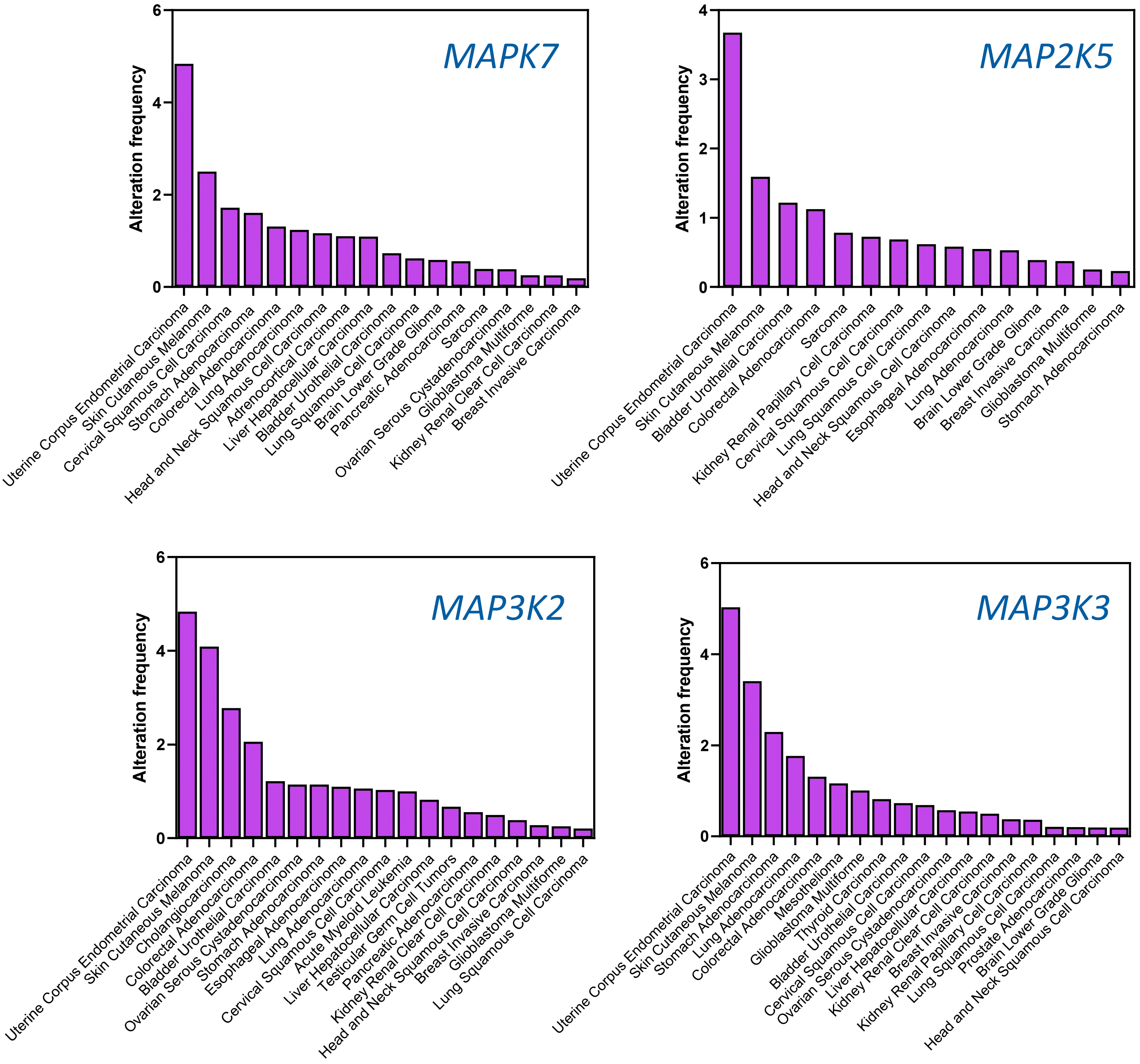


**Figure S1. Mutation frequency of the ERK5 signaling components *MAPK7* (ERK5 *gene*), *MAP2K5 (*MEK5 gene), *MAP3K2* and *MAP3K3* in human cancer.** Cross-cancer alteration summary from 32 studies from TCGA PanCancer Atlas Studies (https://www.cbioportal.org/)

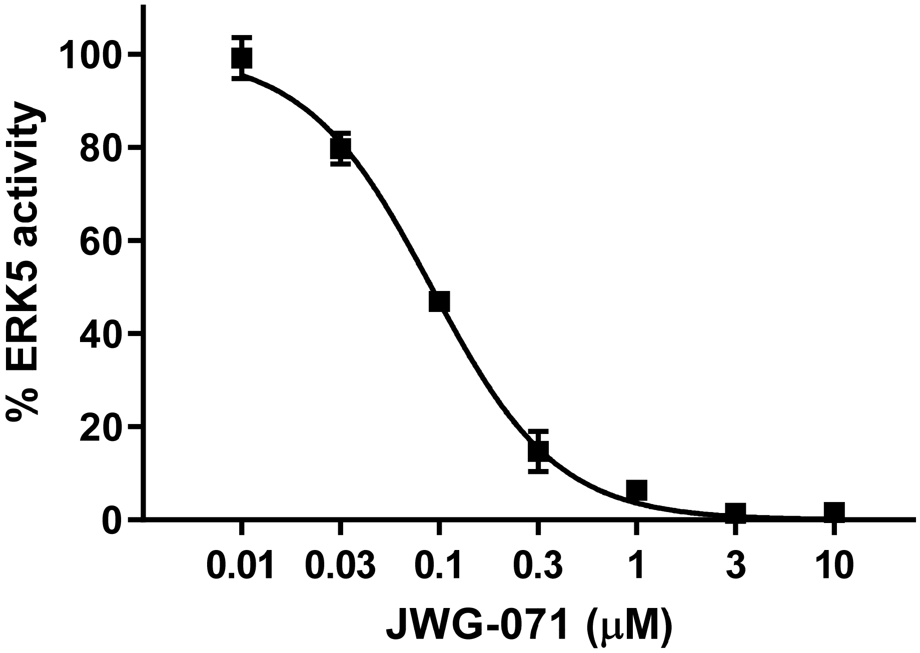


**Figure S2. Radiometric assay of ERK5 kinase activity using the ERK5 specific inhibitor JWG-071.** ERK5 activity was determined using 200 ng of pure recombinant ERK5, 50 μM $^{32}$P-ATP/$Mg^{2+}$ and 200 μM PIMtide as substrates. Assays were carried out for 20 min at 30°C, terminated by applying the reaction mixture onto p81 paper, and the incorporated radioactivity measured by Cherenkov counting. Similar results were obtained in three separate experiments.

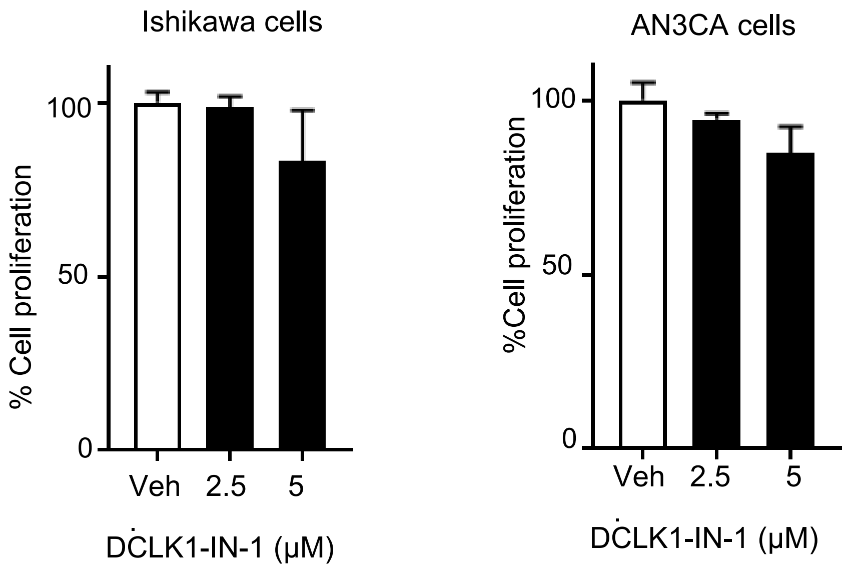


**Figure S3. DCLK1 inhibition does not affect human endometrial cancer cell proliferation.** Ishikawa or AN3CA cells were incubated with the specific DCLK1 inhibitor DCLK1-IN-1 for 48h, and cell proliferation was assessed by MTT assay.

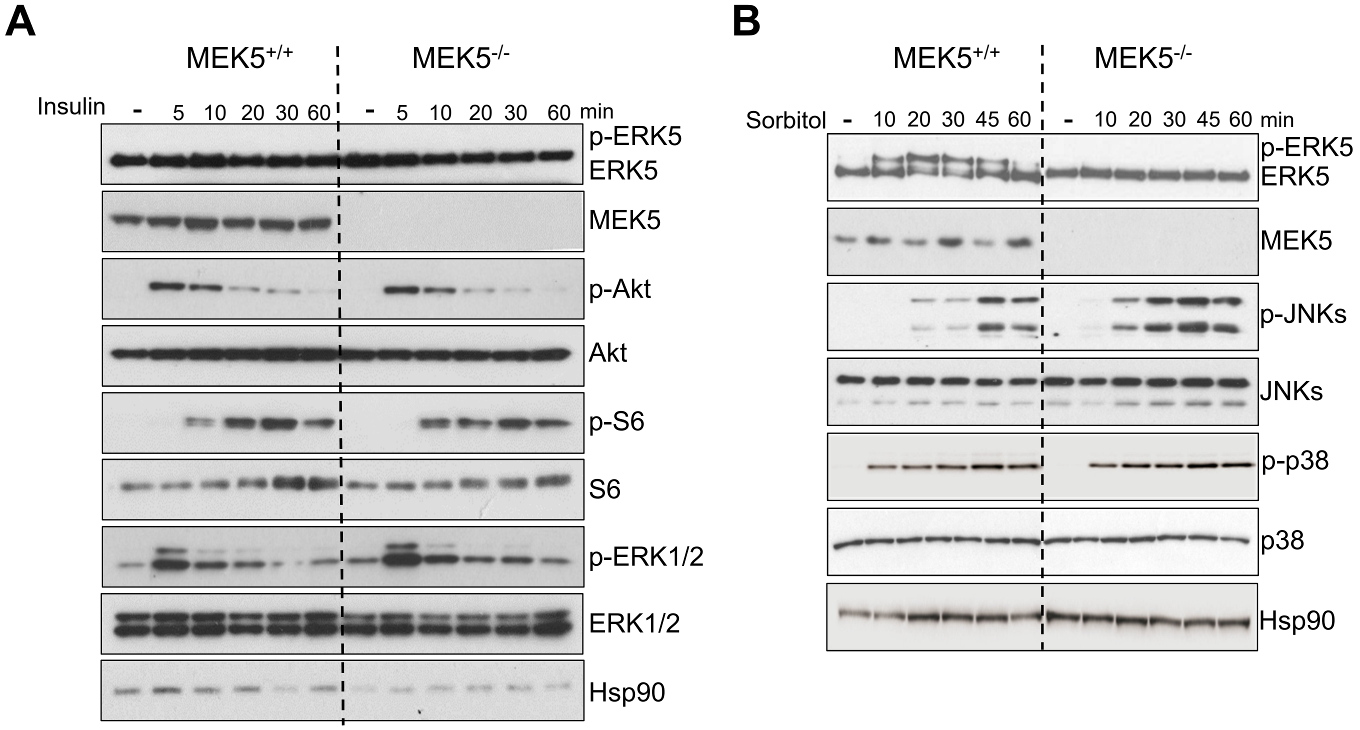


**Figure S4. Effect of sorbitol and insulin stimulation on HeLa MEK5$^{+/+}$ and CRISPR/Cas9 MEK5$^{-/-}$ cells.** Cells were serum starved, previous stimulation with either 50 ng/ml insulin (A) or 0.5 M sorbitol (B) at the indicated times. Expression of the indicated proteins was monitored by immunoblot. Results are representative of two independent experiments.

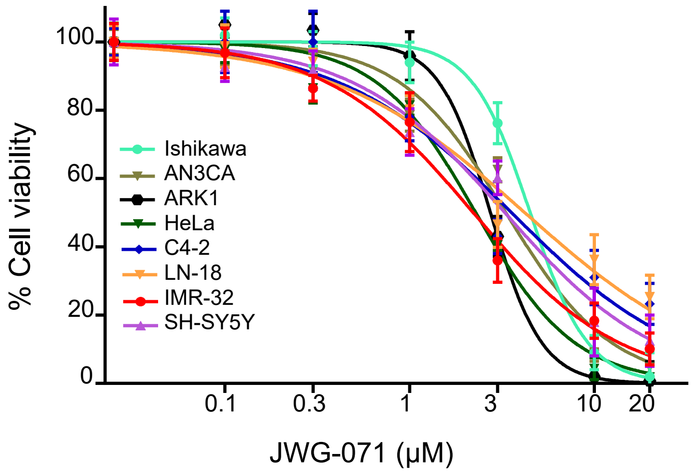


| Cell Line | $IC_{50}$ (μM) |
|---|---|
| Ishikawa (endometrial) | 3.2 ± 0.8 |
| AN3CA (endometrial) | 3.3 ± 0.6 |
| ARK1 (endometrial) | 2.7 ± 0.2 |
| Hela (cervical) | 2.9 ± 0.7 |
| C4-2 (prostate) | 2.2 ± 0.3 |
| LN-18 (glioblastoma) | 2.9 ± 0.4 |
| IMR-32 (neuroblastoma) | 1.9 ± 0.3 |
| SH-SY5Y (neuroblastoma) | 4.2 ± 1.4 |

**Figure S5. ERK5 inhibition (JWG-071) induces cytotoxicity in a panel of human tumor cell lines.** MTT cytotoxicity assay. Cells were incubated with JWG-071 for 48 h. Right table show the corresponding $IC_{50}$ values.

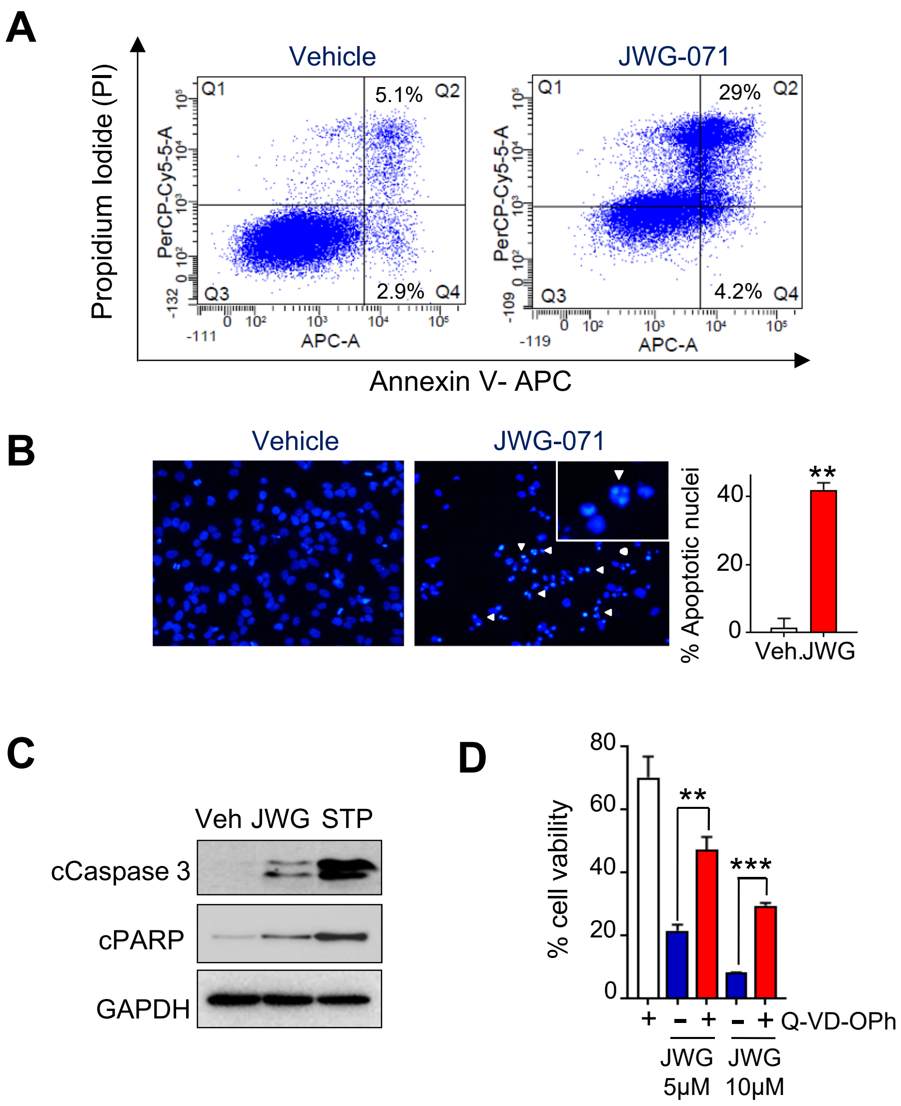


**Figure S6. ERK5 inhibition induces apoptotic cell death in HeLa cells. A,** Percentage of apoptotic cells was determined by Anexin V/PI staining at 48 h following treatment. Representative flow cytometry plots of cells are shown. **B,** Representative images of nuclear morphology (Hoescht 33258) at 48 h post-treatment with 5 μM JWG-071 or vehicle. Arrowheads point condensed or fragmented nuclei. Right histograms, quantification of the results by scoring four representative fields of each condition (n =3). **C,** Immunoblot analysis of cells treated with vehicle or 5 μM JWG-071 (48 h). Staurosporine (STP, 18 h) was used as an apoptosis control. **D,** Pan caspase inhibitor Q-VD-OPh impairs JWG-071-induced cytotoxicity. Cells were preincubated 1 h with 20 mM Q-VD-Oph, before treatment with JWG-071 for 48 h. Cell viability was monitored by MTT assay. Data is presented as the mean of three independent experiments ± SD, each performed in tetraplicates. *, $P < 0.05$; **, $P < 0.005$; ***, $P < 0.001$ (Student's t-test).

**A**

| Histological type | Grade | Staging FIGO | Code |
|---|---|---|---|
| Endometrioid | G1 | Ia | 1 |
| Endometrioid | G1 | Ia | 12 |
| Endometrioid | G1 | Ia | 13 |
| Endometrioid | G2 | Ia | 14 |
| Endometrioid | G2 | Ia | 2 |
| Endometrioid | G3 | IIIc2 | 3 |
| Endometrioid | G3 | Ib | 15 |
| Endometrioid | G2 | II | 4 |
| Endometrioid | G3 | Ib | 5 |
| Endometrioid | G3 | Ia | 16 |
| Endometrioid | G2 | II | 6 |
| Endometrioid | G1 | Ia | 7 |
| Endometrioid | G2 | II | 17 |
| Endometrioid | G2 | IIIc2 | 8 |
| Endometrioid | G3 | Ia | 9 |
| Endometrioid | G3 | IVa | 10 |
| Endometrioid | G2 | Ib | 11 |

**B**

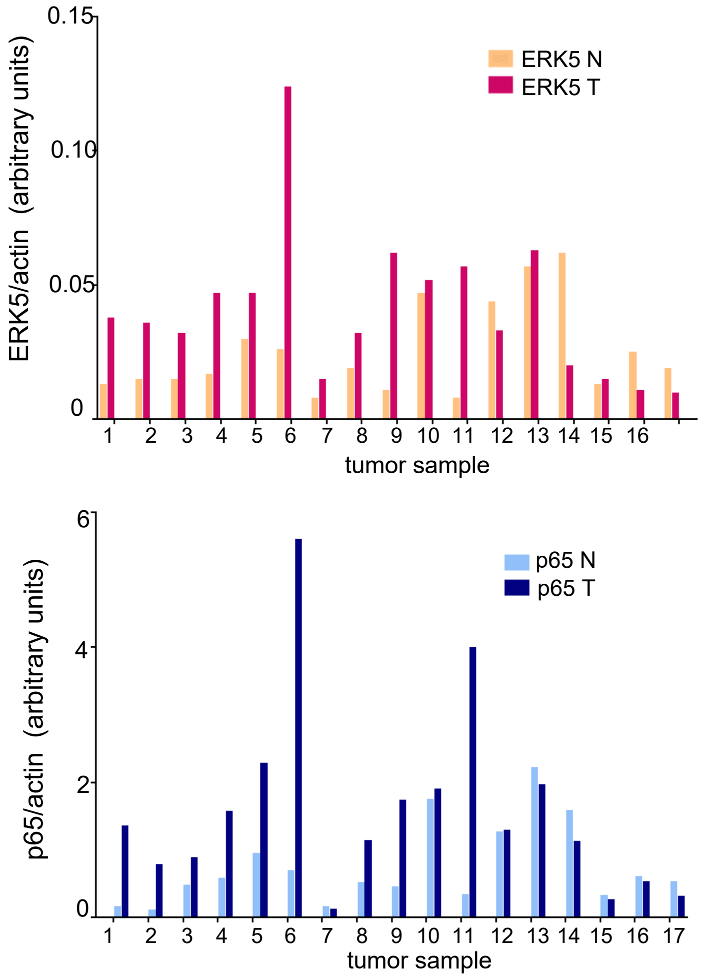


**C**

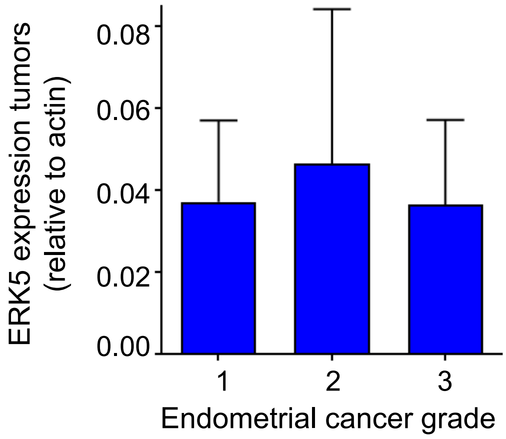


**Figure S7. ERK5 and p65 expression in tumoral and non-tumoral samples from 17 endometrioid cancer patients. A,** Grade and staging (FIGO) of the 17 endometrioid cancer samples**. B,** Quantification of the ERK5 and p65/RELA protein expression, analyzed by immunoblot signal from Figure 4A. Histograms shows the relative ERK5 or p65/RELA values, relative to actin. T, tumoral; N, peritumoral samples. **C,** ERK5 expression on different grades of malignancy.

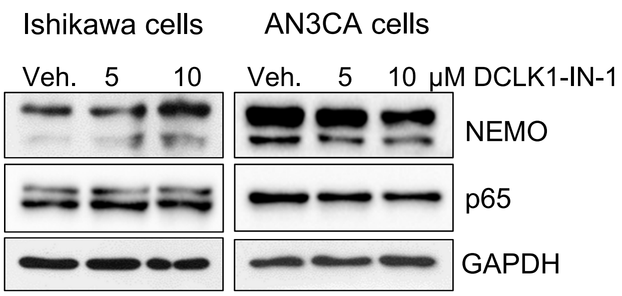


**Figure S8. DCLK1 inhibition does not affect NEMO/IKKγ or p65/RELA protein expression levels.** Ishikawa and AN3CA cells were treated with 5 or 10 µM DCLK1-IN-1 (DCLK1 inhibitor) for 48h. Protein expression was monitored by immunoblot analysis.

A B

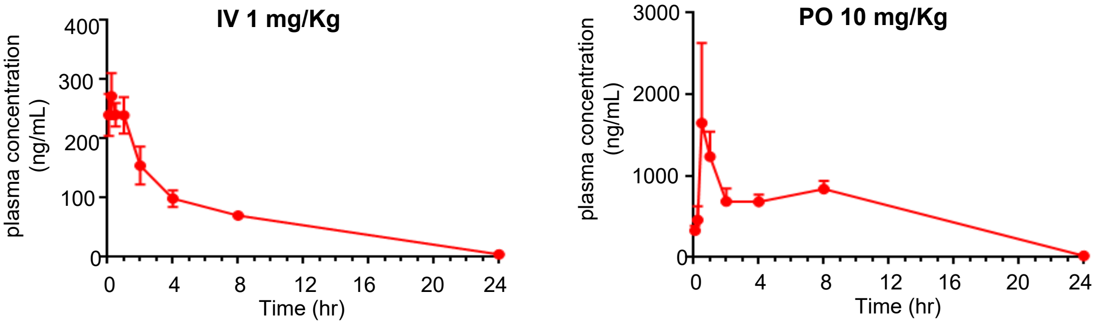


C

| Parameter | Unit | IV | PO |
|---|---|---|---|
| Dose | mg·kg$^{-1}$ | 1 | 10 |
| $T_{max}$ | hr | - | 1.00 |
| $^{a}C_o/C_{max}$ | ng mL$^{-1}$ | 240.07 | 1648.76 |
| $AUC_{last}$ | hr*ng mL$^{-1}$ | 1632.29 | 13778.95 |
| $AUC_{inf}$ | hr*ng mL$^{-1}$ | 1661.28 | 13870.05 |
| $T_{1/2}$ | hr | 4.34 | - |
| CL | mL min$^{-1}$ kg$^{-1}$ | 10.03 | - |
| $V_{ss}$ | L kg$^{-1}$ | 3.35 | - |
| $F^{b}$ | % | - | 84 |

**Figure S9. Pharmacokinetic parameters of JWG-071 (ERK5 inhibidor) in male swiss albino mice. A,** Plot of plasma concentration versus time for intravenous (IV) dosing (1 mg/kg), or **(B)** oral (PO) dosing (10 mg/kg). **C,** Summary of pharmacokinetic parameters. Data in A-B are presented as the mean ±S.D. of measurements from n = 9 independent mice assayed per delivery route.

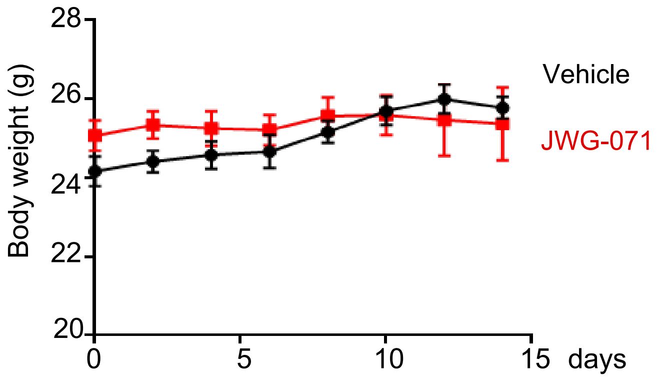


**Figure S10A. Mice body weight variation curve in mouse bearing Ishikawa tumor xenografts treated with vehicle or JWG-071 monotherapy.** Nude mice were daily treated with either vehicle (black line) or with 50 mg/kg JWG-071 (red line). Mice body weight was monitored every two days.

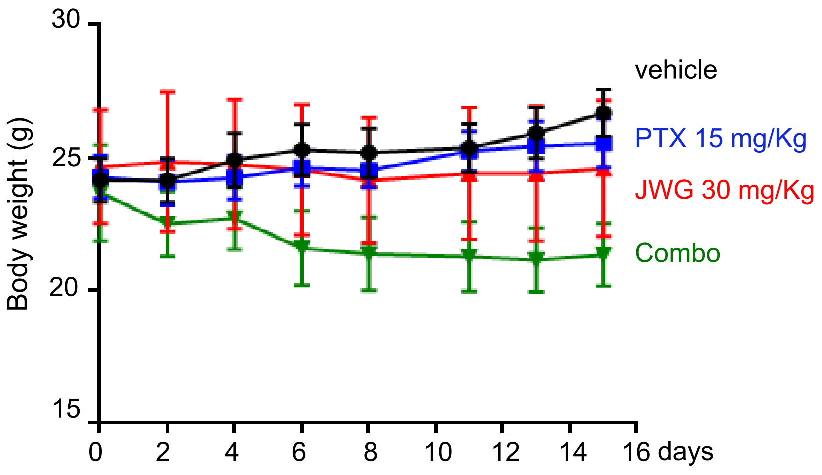


**Figure S10B. Mice body weight variation curve. ERK5i in combination with chemotherapy (paclitaxel).** Nude mice were treated with either vehicle (black line), 15 mg/kg paclitaxel (twice a week, blue line), with 30 mg/kg JWG-071 (daily, red line), or a combination of paclitaxel and JWG-071. Mice body weight was monitored every two days.

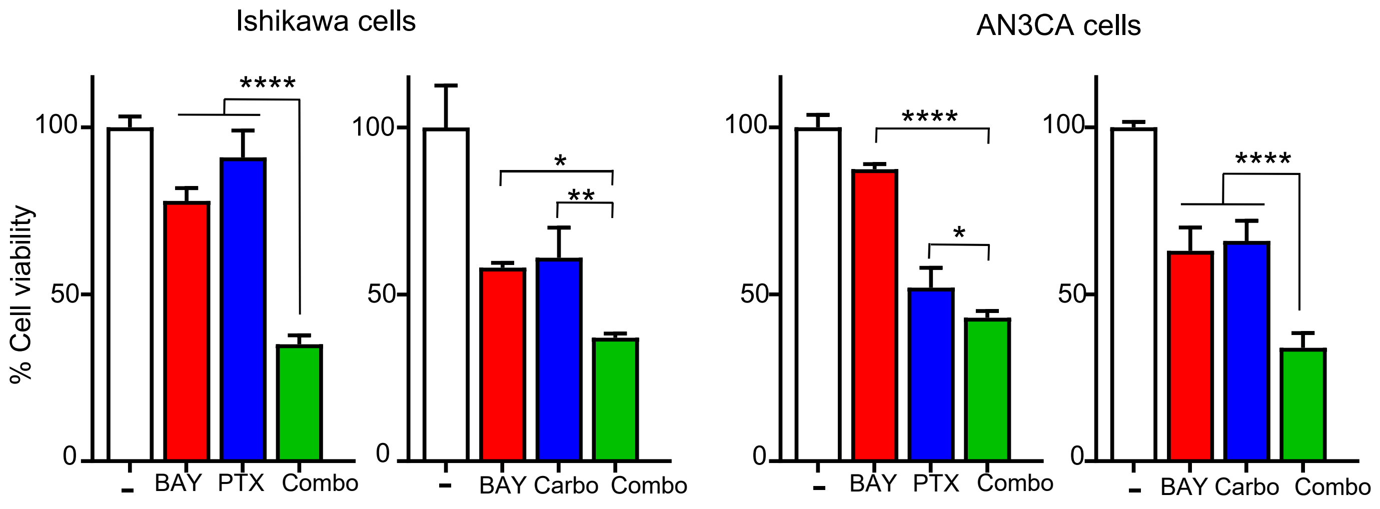


**Ishikawa cells — Paclitaxel (nM) × BAY11-7082 (μM)**

| Paclitaxel (nM) | 0 | 2 | 3 |
|---|---|---|---|
| 0 | 100 | 100 | 78 |
| 0.5 | 100 | 94 | 42 |
| 1 | 91 | 83 | 35 |
| 2 | 45 | 53 | 29 |

**Ishikawa cells — Carboplatin (μM) × BAY11-7082 (μM)**

| Carboplatin (μM) | 0 | 2 | 3 |
|---|---|---|---|
| 0 | 100 | 100 | 58 |
| 20 | 100 | 68 | 56 |
| 30 | 100 | 62 | 56 |
| 40 | 61 | 42 | 37 |

**AN3CA cells — Paclitaxel (nM) × BAY11-7082 (μM)**

| Paclitaxel (nM) | 0 | 0.5 | 1 |
|---|---|---|---|
| 0 | 100 | 92 | 81 |
| 0.25 | 59 | 68 | 62 |
| 0.5 | 52 | 60 | 44 |
| 0.7 | 40 | 36 | 29 |

**AN3CA cells — Carboplatin (μM) × BAY11-7082 (μM)**

| Carboplatin (μM) | 0 | 0.5 | 1 |
|---|---|---|---|
| 0 | 100 | 98 | 73 |
| 100 | 86 | 80 | 58 |
| 120 | 78 | 73 | 44 |
| 140 | 66 | 70 | 34 |

**Figure S11. The NF-κB inhibitor BAY11-7082 sensitizes endometrioid cancer cells to paclitaxel and carboplatin toxicity.** Cells were treated with the indicated concentrations of BAY11-7082, carboplatin (Carbo) or paclitaxel (PTX) for 48 h, and cell viability was determined by MTT assay. Lower panels: Heat-map analysis of the viability values obtained for the tested inhibitors.

Fig. 2F Tumor growth curves

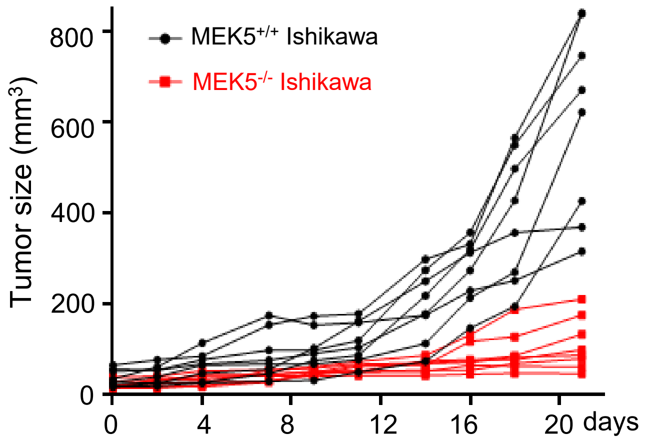


Fig. 6A Tumor growth curves

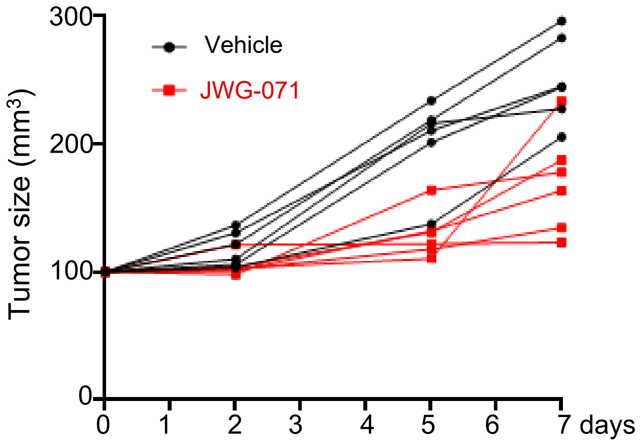


Fig. 6C Tumor growth curves

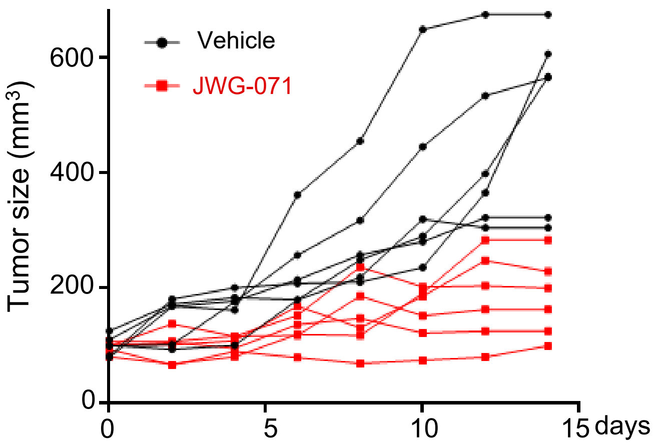


Fig. 7D Tumor growth curves

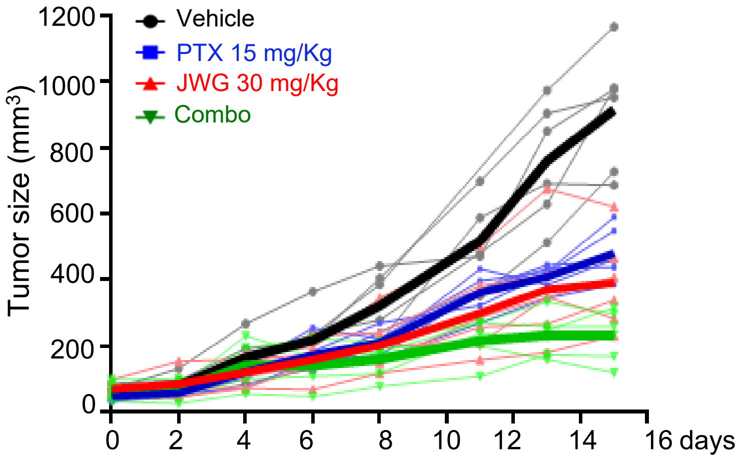


**Figure S12. Individual tumor growth curves corresponding to the *in vivo* experiment in mice.**

| Primary antibody | Produced by | Supplied by | Concentration |
|---|---|---|---|
| β-Actin | Mouse | Santa Cruz, sc-47778 | WB 1:5,000 |
| AKT | Rabbit | Cell Signalling, 9272 | WB 1:5,000 |
| Phospho-AKT (S473) | Rabbit | Cell Signalling, 9271 | WB 1:1,000 |
| Bak | Rabbit | Cell Signalling, 12105 | WB 1:1,000 |
| Bax | Mouse | BD Biosciences, 556467 | WB 1:1,000 |
| Bcl-2 | Mouse | Cell Signalling, 15071 | WB 1:1,000 |
| Bim | Rabbit | Cell Signalling, 2933 | WB 1:1,000 |
| BrdU | Mouse | BD Pharmingen | IF 1:200 |
| Cleaved caspase-3 | Rabbit | Cell Signalling, 9661 | WB 1:500<br>IHC 1:200 |
| Cleaved PARP | Rabbit | Cell Signalling, 5625 | WB 1:1,000 |
| CREB-1 | Mouse | Santa Cruz, sc-186 | WB 1:200 |
| ERK1/2 | Rabbit | Cell Signalling, 4695 | WB 1:8,000 |
| Phospho-ERK1/2 | Rabbit | Cell Signalling, 4376 | WB 1:5,000 |
| ERK5 | Rabbit | Cell Signalling, 3372 | WB 1:1,000 |
| GAPDH | Mouse | Invitrogen, AM4300 | WB 1:100,000 |
| GST | Rabbit | Santa Cruz, sc-459 | IC 1:500 |
| Hsp90-β | Rabbit | Invitrogen, PA3-012 | WB 1:10,000 |
| IKK-α | Mouse | Cell Signalling, 11930 | WB 1:1,000 |
| IKK-β | Rabbit | Cell Signalling, 8943P | WB 1:1,000 |
| IKK-$\gamma$/NEMO | Mouse | Cell Signalling, 2695 | WB 1:1,000 |
| IKB-α | Mouse | Cell Signalling, 4814 | WB 1:1,000 |
| JNK | Rabbit | Cell Signalling, 9252 | WB 1:1,000 |
| Phospho-JNK | Rabbit | Cell Signalling, 4668 | WB 1:1,000 |
| Ki67 | Rabbit | Cell Signalling, 5365 | IHC 1:600 |
| MEK5 (E-3) | Mouse | Santa Cruz, sc-365198 | WB 1:250 |
| p65 | Rabbit | Cell Signalling, 8242 | WB 1:1,000 |
| Phospho-p65 | Rabbit | Cell Signalling, 3033 | WB 1:1,000 |
| S6 | Rabbit | Cell Signalling, 2217 | WB 1:20,000 |
| Phospho-S6 | Rabbit | Cell Signalling, 4858 | WB 1:40,000 |

**Supplementary Table1.** List of antibodies used.
